# Hamsters are a model for post-COVID-19 alveolar regeneration mechanisms: an opportunity to understand post-acute sequelae of SARS-CoV-2

**DOI:** 10.1101/2022.11.17.515635

**Authors:** Laura Heydemann, Małgorzata Ciurkiewicz, Georg Beythien, Kathrin Becker, Klaus Schughart, Stephanie Stanelle-Bertram, Berfin Schaumburg, Nancy Mounogou-Kouassi, Sebastian Beck, Martin Zickler, Mark Kühnel, Gülsah Gabriel, Andreas Beineke, Wolfgang Baumgärtner, Federico Armando

**Author notes:** **Correspondence** and requests for materials should be addressed to Wolfgang Baumgärtner. These authors contributed equally as co-first authors (Laura Heydemann, Małgorzata Ciurkiewicz). These authors contributed equally as co-last authors (Wolfgang Baumgärtner, Federico Armando).

## Abstract

A relevant number of coronavirus disease 2019 (COVID-19) survivors suffers from post-acute sequelae of severe acute respiratory syndrome coronavirus 2 (PASC). Current evidence suggests a dysregulated alveolar regeneration in COVID-19 as a possible explanation for respiratory PASC symptoms, a phenomenon which deserves further investigation in a suitable animal model. This study investigates morphological, phenotypical and transcriptomic features of alveolar regeneration in SARS-CoV-2 infected Syrian golden hamsters. We demonstrate that CK8^+^ alveolar differentiation intermediate (ADI) cells occur following SARS-CoV-2-induced diffuse alveolar damage. A subset of ADI cells shows nuclear accumulation of TP53 at 6- and 14-days post infection (dpi), indicating a prolonged arrest in the ADI state. Transcriptome data show the expression of gene signatures driving ADI cell senescence, epithelial-mesenchymal transition, and angiogenesis. Moreover, we show that multipotent CK14^+^ airway basal cell progenitors migrate out of terminal bronchioles, aiding alveolar regeneration. At 14 dpi, presence of ADI cells, peribronchiolar proliferates, M2-type macrophages, and sub-pleural fibrosis is observed, indicating incomplete alveolar restoration. The results demonstrate that the hamster model reliably phenocopies indicators of a dysregulated alveolar regeneration of COVID-19 patients. The results provide important information on a translational COVID-19 model, which is crucial for its application in future research addressing pathomechanisms of PASC and in testing of prophylactic and therapeutic approaches for this syndrome.

## Introduction

Severe acute respiratory syndrome coronavirus 2 (SARS-CoV-2), caused over 600 million infections and over 6.5 million fatal outcomes to this day (October 2022, WHO). Patients surviving acute COVID-19 are at risk to develop post-acute sequelae of SARS-CoV-2 (PASC) ^1-3^. PASC occurs in 3-11.7% of infected individuals and is characterized by symptoms such as fatigue, headache, cognitive dysfunction, altered smell and taste, shortness of breath, and dyspnea, occurring >12 weeks after acute virus infection ^4,5^. Of note, among patients with severe disease requiring hospitalization, shortness of breath or dyspnea are reported with a much higher frequency (in up to 49% and 23.3% of cases, respectively) 8-10 months after acute disease ^6,7^. The pathomorphological correlates and mechanisms responsible for respiratory PASC are still not fully understood. Impairment of gas exchange capacity due to an incomplete or protracted regeneration of alveoli and lung fibrosis represent potential pathomechanisms ^8-10^. SARS-CoV-2 infection of the lung causes diffuse alveolar damage (DAD), characterized by necrosis of alveolar epithelial type 1 and 2 (AT1 and AT2) cells, fibrin exudation and edema, followed by alveolar epithelial hyperplasia in later stages ^11,12^. The healing of damaged alveoli and recovery of gas exchange capacity requires the presence of progenitor cells that are able to regenerate lost AT1 cells. For a long time, it was assumed that AT1 cells are regenerated solely by proliferating and trans-differentiating AT2 cells. However, recent advances in mouse models of lung injury have shown that different airway progenitor cell types expand and mobilize to repair alveolar structures ^13-16^. AT2 cells are mainly responsible for AT1 cell regeneration in homeostatic turnover and following mild injury, while airway progenitors are recruited after severe injury with marked AT1 cell loss ^15,17^. The differentiation into mature AT1 cells features an intermediate step, the so-called *alveolar differentiation intermediate* (ADI) cell, first described to occur during AT2 to AT1 cell trans-differentiation ^18-21^. ADI cells in mice are characterized by cytokeratin 8 (CK8) expression, a polygonal to elongated morphology, NFκB and TP53 activation and upregulation of genes involved in epithelial– mesenchymal transition (EMT), HIF-1α pathway, and cell cycle exit ^8,19^. ADI cells have been observed in various lung injury models, e.g. bleomycin injury, neonatal hypoxia and hyperoxia, LPS injury and Influenza A virus infection ^18-20^. In homeostatic turnover and mild injury, these cells occur only transiently and differentiate into mature AT1 cells eventually, thereby restoring normal alveolar structure and function ^19,21,22^. However, a pathological accumulation of ADI cellshas been observed in idiopathic pulmonary fibrosis (IPF) in humans and a mouse model for progressive fibrosis, suggesting that a blockage during trans-differentiation of ADI to AT1 cells could represent a potential regenerative defect in these conditions ^8,18,19,21,23^. Recently, high numbers of ADI cells have also been demonstrated in lungs of COVID-19 patients. It has been postulated that persistence of these cells could be responsible for unremitting hypoxemia, edema, ventilator dependence and the fatal outcome in protracted ARDS as well as the subsequent development of fibrosis in PASC ^8,9,24,25^. Since serial samples are rarely available in human observational studies, the fate of COVID-19 associated ADI cells remains elusive. Addressing this open question is of paramount importance to obtain a deeper understanding of the factors that contribute to the protracted recovery from COVID-19 facilitating the development of rational therapeutic approaches in the field of lung regenerative medicine. The development of a precise working hypothesis and subsequent preclinical testing of therapeutic options requires the study of sequential phases of SARS-CoV-2 infection in appropriate animal models. Among the susceptible small animal species, Syrian golden hamsters (*Mesocricetus auratus*) are well suited to study regenerative responses. They develop a distinct, but transient and non-lethal disease, in contrast to other models such as transgenic mice or ferrets ^26-30^. The regeneration of lung epithelia following SARS-CoV-2 infection has not been characterized in detail in this important animal model yet. To obtain further insights in the regenerative processes following SARS-CoV-2 infection we characterized the proliferating epithelial cells within the lung of infected hamsters in the acute and sub-acute phase of the infection until 14 days post infection. Our study shows that CK8^+^ADI cells and multipotent CK14^+^ airway basal cells participate in alveolar regeneration and that persistence of ADI cells at 14 dpi is associated with fibrosis in SARS-CoV-2 infected hamsters. In addition our study provides a hamster-specific marker gene lists for different alveolar cell populations, including AT1, ADI and AT2 cells. Altogether, the results provide important information on a translational COVID-19 model, which is crucial for its application in future research addressing pathomechanisms of PASC and for testing of prophylactic and therapeutic approaches for this syndrome.

## Results

### 1. SARS-CoV-2 induced epithelial proliferative responses and inflammation persist beyond virus clearance

First, successful infection was confirmed by immunohistochemistry for SARS-CoV-2 nucleoprotein (NP) antigen in lung tissue. Viral antigen was found in alveolar and bronchial epithelia as well as in macrophages (**Fig. 1A**), as described previously ^31^. Quantification of immunolabeled cells in whole lung sections peaked at 3 dpi, followed by a sharp decline at 6 dpi and virus clearance at 14 dpi (**Fig. 1A**). No SARS-CoV-2^+^ cells were detected in mock-infected animals at any time point. Histologically, SARS-CoV-2 infected animals showed a marked, transient, broncho-interstitial pneumonia, as described previously ^31-34^. The lesions were characterized by DAD with epithelial cell degeneration and necrosis, sloughing of alveolar cells, fibrin exudation and heterophilic and histiocytic infiltrates. Some mock-infected animals showed small foci of mild, multifocal, interstitial inflammation composed of heterophils and macrophages, particularly at 1 dpi. The extent of inflammation in SARS-CoV-2 and mock infected animals was quantified in total lung sections using whole slide digital image analysis of Iba-1 immunolabeling. The number of Iba-1^+^ cells was significantly higher in SARS-CoV-2 infected animals compared to the mock group at 1, 3, and 6, with a notable peak at 6 dpi (**Fig. 1B**).

**Figure 1.**
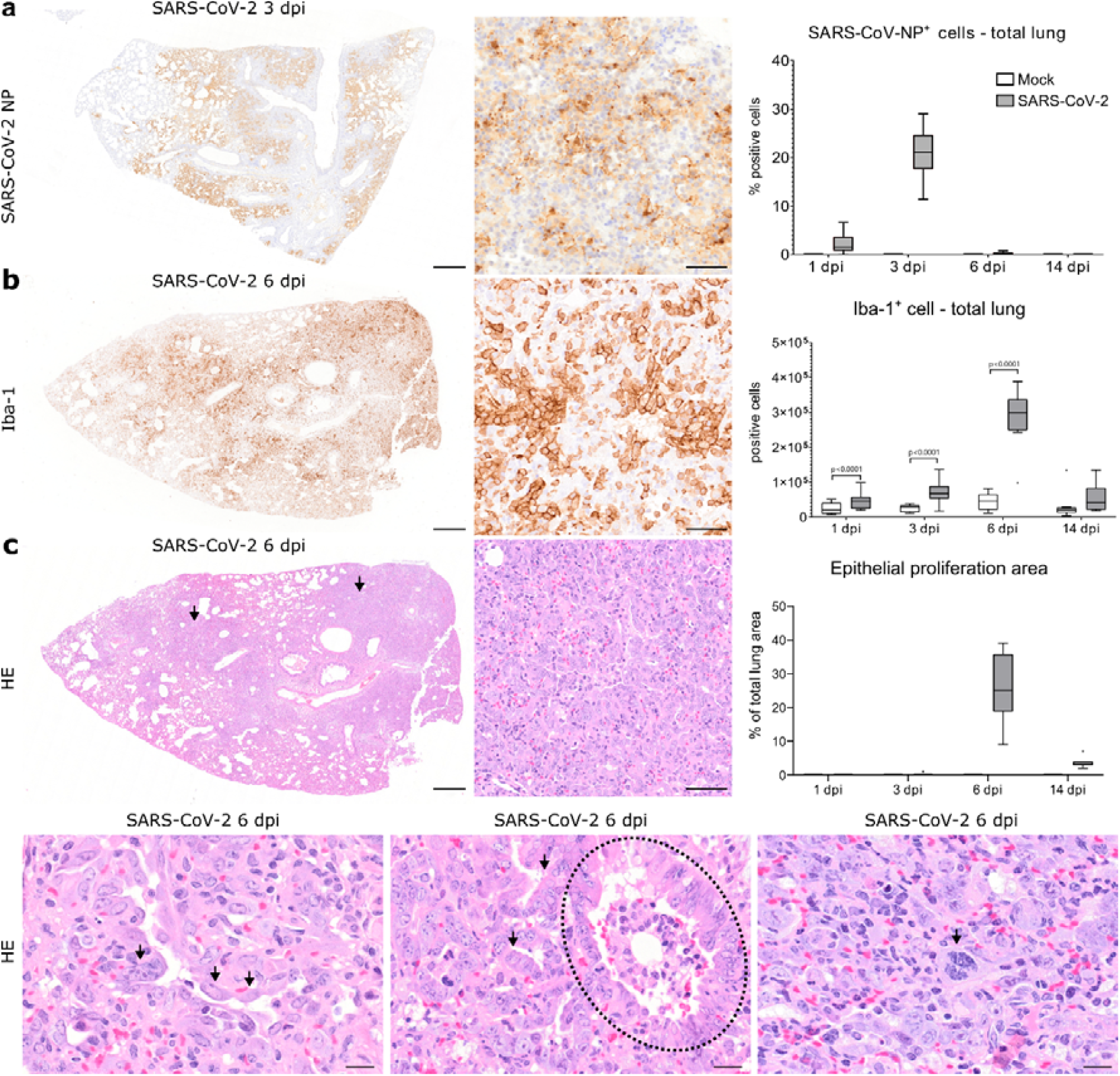
SARS-CoV-2 infection causes a marked epithelial proliferative response in the hamster lung. A Representative images showing SARS-CoV-2 nucleoprotein (NP) immunolabeling in one right lung lobe of an infected hamster at 3 days post infection (dpi). The left panel shows an overview of one right lung lobe and the central panel displays a higher magnification of viral antigen (brown signal) in the alveoli. Quantification of SARS-CoV-2 NP^+^ cells is shown in the right panel. **B** Representative images showing ionized calcium-binding adapter molecule 1 (Iba-1) immunolabeling in one right lung lobe of an infected hamster at 6 dpi. The left panel shows an overview of one right lung lobe and the central panel display a higher magnification of macrophages/histiocytic cells (brown signal) in the affected alveoli. Quantification of Iba-1^+^ macrophages/histiocytes is shown in the right panel. **C** Representative images showing histopathological lesions in a lung lobe of a SARS-CoV-2 infected hamster at 6 dpi. The top left panel shows an overview of one right lung lobe displaying large areas of alveolar consolidation (arrows). The top central panel shows a higher magnification of an affected region, which shows a prominent epithelial proliferation. The quantification of epithelial proliferation is reported in the top right panel. The percentage of affected area relative to total lung area is given. The bottom left panel shows strings of plump polygonal or elongated cells lining alveolar septa (arrows). The bottom central panel shows proliferation of cuboidal airway epithelial cells forming ribbons and tubules (arrows) surrounding terminal bronchioles (dotted line). The bottom right panel shows that within alveolar proliferation foci, there are cells displaying karyomegaly and atypical mitotic figures (arrow). Data are shown as box and whisker plots. Data from Iba-1 quantification was tested by two-tailed Mann-Whitney-U test. A p-value of ≤0.05 was considered significant. N□=□10 animals/group for mock and SARS-CoV-2 respectively. For quantifications, 1 longitudinal section containing all right lung lobes were evaluated. Source data will be provided as a source data file. Scale bars: 500 µm (overviews in a-c), 50 µm (high magnifications in a-c), 20 µm (high magnifications in lower panel in c).

The inflammatory lesions in SARS-CoV-2 infected hamsters were accompanied by a prominent epithelial proliferation (**Fig. 1C**). At 3 dpi, small foci of hyperplastic epithelial cells were observed within alveoli in single animals, affecting up to 1.3% of the examined lung area (**Fig. 1C**). At 6 dpi, large areas of prominent epithelial cell proliferation were found in all infected animals, affecting 9.3% to 39.3% of the examined lung area (**Fig. 1C**). Proliferating epithelial cells within the alveoli were characterized by variable morphologies, including a round cell shape typical of AT2 cells and a more polygonal to sometimes elongated shape resembling ADI cells. Surrounding terminal bronchioles, a proliferation of cuboidal airway epithelial cells forming pods, ribbons and tubules was observed. In the periphery, these peri-bronchiolar proliferates merged with areas of alveolar epithelial hyperplasia, showing a transition from a cuboidal to a polygonal shape (**Fig. 1C**). Many cells showed atypical features such as cyto- and karyomegaly, bizarrely shaped and euchromatic nuclei, as well as abundant, partly atypical, mitotic figures (**Fig. 1C**). At 14 dpi, multifocal areas of epithelial proliferates were still observed, affecting 2.1% to 7.2% of the examined lung area, often around terminal airways (**Fig. 1C**). In addition, a majority of animals (7 out of 9) showed foci of sub-pleural fibrosis.

In summary, SARS-CoV-2-infected hamsters showed a prominent and heterogeneous epithelial proliferative response that was still recognizable at 14 dpi, beyond virus clearance. Next, we wanted to demonstrate that alveolar AT2 cells proliferate, mobilize and differentiate into AT1 cells through the ADI cell state and that airway-derived progenitors participate in alveolar regeneration, possibly through a transitional AT2 or ADI cell state, in the Syrian golden hamster.

### 2. CK8^+^ ADI cells frequently express TP53 and persist until 14 dpi following SARS-CoV-2 induced DAD in hamsters

ADI cells are reported to originate from AT2 and/or a particular subset of club cells expressing MHC-II ^19^. The AT2 to ADI cell trans-differentiation process is characterized by gradual down-regulation of AT2 cell markers, expression of CK8 and cell cycle exit markers, as well as a morphologic transition from a round to a polygonal to elongated shape ^19,20^. In the following, we focused on the first part of this AT2-ADI-AT1 trajectory (**Fig. 2A**).

**Figure 2:**
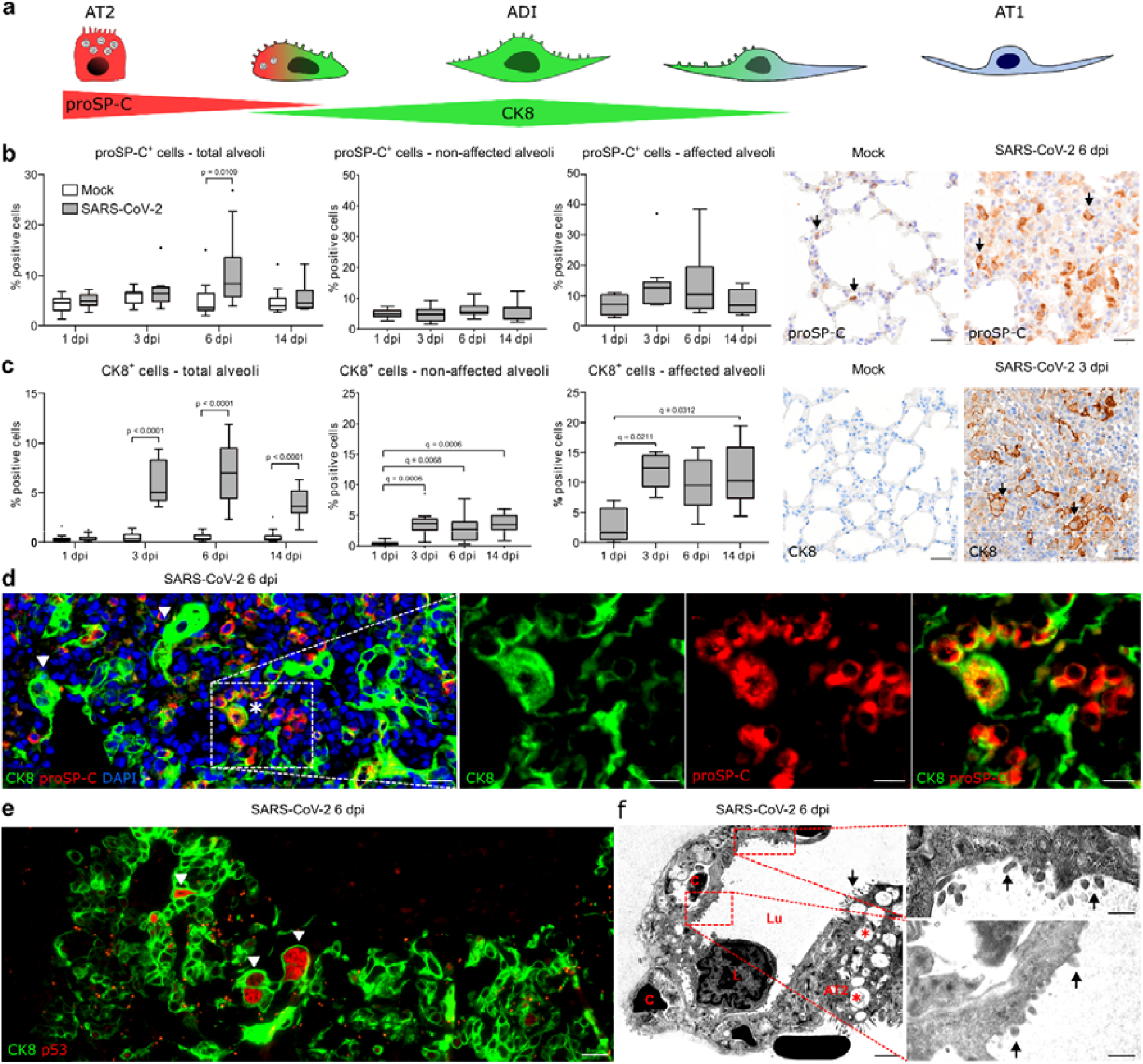
Alveolar differentiation intermediate (ADI) cells in SARS-CoV-2 infected hamsters. A Schematic illustration of the trans-differentiation process from alveolar pneumocytes type 2 (AT2) to alveolar pneumocytes type 1 (AT1), as demonstrated below in the hamster. The AT2 to ADI cell trans-differentiation process is characterized by a decrease of pro surfactant protein-C (proSP-C) expression, increase of cytokeratin 8 (CK8) expression as well as a morphologic transition from a round to polygonal to elongated shape (see D). The ultrastructural hallmark of the ADI to AT1 trans-differentiation is the presence of cells with an elongated AT1 morphology and AT2 features, such as apical microvilli (see F). **B, C** Quantification of proSP-C^+^ AT2 cells (B) and CK8^+^ ADI cells (C) within total alveoli, non-affected alveoli, and affected alveoli as well as representative pictures of immunolabelling (brown signal, arrows) in the alveoli of mock and SARS-CoV-2 infected hamsters. **D** Representative double immunofluorescence image of an alveolar proliferation focus in a SARS-CoV-2 infected hamster at 6 dpi. Cells are labelled with CK8 (green) and proSP-C (red). An overview and higher magnification of the area delineated by a rectangle are shown. There are numerous proSP-C^-^CK8^+^ADI cells, some showing hypertrophy and elongated cytoplasmic processed (arrowheads) and single proSP-C^+^CK8^+^ cells (asterisk) with a round morphology. **E** Representative double immunofluorescence of an alveolar proliferation focus in a SARS-CoV-2 infected hamster at 6 dpi. Cells are labelled with CK8 (green) and cell cycle exit marker TP53 (red). The arrowheads shows polygonal, large, bizarre TP53^+^ ADI cells. **F** Representative transmission electron microscopy (TEM) micrograph showing alveoli of a SARS-CoV-2 infected hamster at 6 dpi. A basement membrane separates AT1 cells from the endothelial cells lining capillary spaces (C) containing erythrocytes. A leukocyte (L) as well as an AT2 cell (AT2) with apical microvilli (arrow) and numerous intracytoplasmic multi-lamellar bodies (red asterisks) are also seen. Red boxes and high magnification show cells with flattened and elongated morphology of AT1 cells, with characteristics of AT2 cells, such as microvilli (arrows). Quantification data are shown as box and whisker plots. Statistical analysis was performed by two-tailed Mann-Whitney-U test. For multiple comparisons between time points, a Benjamini– Hochberg correction was applied. P- and q-values ≤0.05 were considered significant. ND=D10 animals/group for mock and SARS-CoV-2 respectively. For quantifications, 1 longitudinal section containing all right lung lobes were evaluated. Source data will be provided as a source data file. Scale bars: 25 µm (b, c), 20 µm (overview in d, e), 10 µm (high magnification in d), 2000 nm (low magnification in f), 500 nm (high magnification in f).

First, we detected proSP-C^+^ AT2 and CK8^+^ ADI cells using immunohistochemistry. Quantification was performed within total alveoli first, followed by a separate analysis in areas showing inflammation and/or epithelial proliferation (termed “affected alveoli”) and histologically unremarkable alveoli (termed “non-affected alveoli”). proSP-C expression was detected in cells with a round shape lining alveolar septa. In mock-infected animals, the number of proSP-C^+^ cells was constant at all investigated time-points (**Fig. 2B**). In SARS-CoV-2 infected animals, the total number of proSP-C^+^ cells increased significantly at 6 dpi, which was caused by an increase within affected alveoli. proSP-C^+^ cells were found in small groups within inflammatory foci (**Fig. 2B**). Scattered proSP-C^+^ cells were observed in close proximity of terminal bronchioles. Interestingly, the majority of cells within the epithelial proliferates at 6 dpi were proSP-C^-^.

CK8 was ubiquitously expressed in the apical cytoplasm of luminal cells within bronchi, bronchioles and terminal bronchioles in all animals. In the alveoli of mock-infected animals, rare elongated CK8^+^ cells were observed, making up less than 1% of total alveolar cells. In SARS-CoV-2 infected animals however, CK8 was abundantly expressed within the epithelial proliferative foci at 3, 6 and 14 dpi and the number of CK8^+^ cells in total alveoli was significantly increased compared to the mock group (**Fig. 2C**). Importantly, increased numbers of CK8^+^ cells were detected within affected and non-affected alveoli. Within affected alveoli, the relative numbers of CK8^+^ cells remained constantly elevated throughout the investigation period. CK8^+^ cells displayed a variable cell morphology including round, polygonal, as well as elongated shapes with thin cytoplasmic processes (**Fig. 2C**).

Double-labeling for proSP-C and CK8 demonstrated AT2 to ADI cell transition. At 3 dpi, numerous proSP-C^+^CK8^-^ cells and rare proSP-C^+^CK8^+^ cells with a round AT2 cell morphology were observed within affected alveoli (Supplem. Fig. 1A), whereas proSP-C^-^CK8^+^ elongated cells were very rare. At 6 dpi, affected alveoli contained occasional proSP-C^+^CK8^-^ and proSPC^+^CK8^+^ round cells (**Fig. 2D**). These cells were intermingled with high numbers of proSP-C^-^CK8^+^cells, which showed various morphologies ranging from round AT2-type to polygonal ADI-type cells as well as bizarre, irregularly shaped cells with karyomegaly. Moreover, elongated proSPC^-^ CK8^+^ cells with AT1-type morphology were occasionally observed (Supplem. Fig. 1B). At 14 dpi, numerous proSPC^+^CK8^+^ cells with AT2 morphology as well as occasional proSPC^-^CK8^+^ polygonal cells were still detected in alveoli, including morphologically non-affected alveoli (Supplem. Fig. 1C).

Once AT2 cells enter the ADI state, they exit the cell cycle to allow AT1 trans-differentiation ^8,21^. At 6 and 14 dpi, CK8^+^ cells in SARS-CoV-2 infected animals expressed nuclear TP53, indicative of cell cycle arrest and DNA repair (**Fig. 2E)** (Supplem. Fig. 2A-B). TP53 expression was particularly frequent in polygonal, large, bizarre, occasionally bi-nucleated cells (Supplem. Fig. 2A-B). Of note, no TP53 expression was observed in the rare CK8^+^ cells in the alveoli of mock-infected animals. Our findings demonstrated that the transition between AT2 and ADI cells in SARS-CoV-2 infected hamsters features: i) transient co-expression of proSP-C and CK8, ii) changes in cell morphology from round to elongated as well as iii) expression of cell cycle arrest markers.

In functional regeneration, ADI cells transdifferentiate into mature AT1, which assume an elongated morphology with thin cytoplasmic processes required for adequate gas exchange. In the following, we focused on the last part of this AT2-ADI-AT1 trajectory. Double-labeling with AT1 cell markers described for human and mouse (AGER, AQP5, PDPN) was not possible since the tested antibodies failed to specifically label AT1 cells in the hamster (*data not shown*). For this reason, we performed transmission electron microscopy to demonstrate ADI to AT1 cell transition. In normal conditions, AT1 cells are characterized by a flattened morphology with slender processes containing a moderately electron-dense, organelle-poor cytoplasm and a round to oval nucleus with a moderate amount of peripheral heterochromatin^35,36^. AT2 cells are characterized by a round morphology, an apico-basal polarity and a moderately electron-dense cytoplasm rich in rough endoplasmic reticulum and free ribosomes. In addition, AT2 cells possess apical microvilli as well as membrane-bound vesicles containing multiple concentric membrane layers (multi-lamellar bodies)^35,36^. In SARS-CoV-2 infected animals, proliferative foci at 6 dpi contained numerous AT2 cells (Supplem. Fig. 3A) as well as numerous hypertrophic epithelial cells with a variable cell morphology resembling ADI cells (Supplem. Fig. 3B; Supplem. Fig. 4 A-B). Most importantly, cells sharing AT1 and AT2 cell features were observed in the alveolar lining at the edges of proliferative foci. The cells showed the flattened and elongated morphology of AT1 cells, but also characteristics of AT2 cells, such as microvilli on the cell surface (**Fig. 2F;** Supplem. Fig. 3C-D). Similar ultrastructural findings were present in COVID-19 patients ^37^. The present findings demonstrate that the last part of the AT2-ADI-AT1 trajectory also occurs in SARS-CoV-2 infected hamsters.

Finally, we sought to confirm that the ADI cells detected in the hamster share features with ADI cells in COVID-19 patients. For this, we used lung samples obtained from three patients with lethal COVID-19 ARDS. In addition, a fourth lung sample obtained from a lobectomy of a non-COVID case was used. Histologically, the lungs from all lethal COVID-19 ARDS cases showed features of moderate to severe, acute DAD, characterized by necrosis and sloughing of alveolar cells, fibrin exudation, hyaline membranes, alveolar edema and mild to moderate neutrophilic infiltrates (**Fig. 3A)**. In the non-COVID-19 sample, a suppurative bronchopneumonia was diagnosed, characterized by neutrophilic and histiocytic infiltrates in bronchioles and alveolar lumina (**Fig. 3A)**. Immunolabeling showed the presence of round proSP-C^+^CK8^+^ cells and polygonal to elongated proSP-C^-^CK8^+^ cells, representing the different stages of ADI cells, in all lethal COVID-19 ARDS samples as well as the non-COVID-19 bronchopneumonia sample (**Fig. 3B)**. Interestingly, CK8^+^ ADI cells expressing TP53 were only detected the three lethal COVID-19 ARDS samples, while no TP53 co-expression was detected in the ADI cells of the non-COVID-19 case (**Fig. 3C)**.

**Figure 3:**
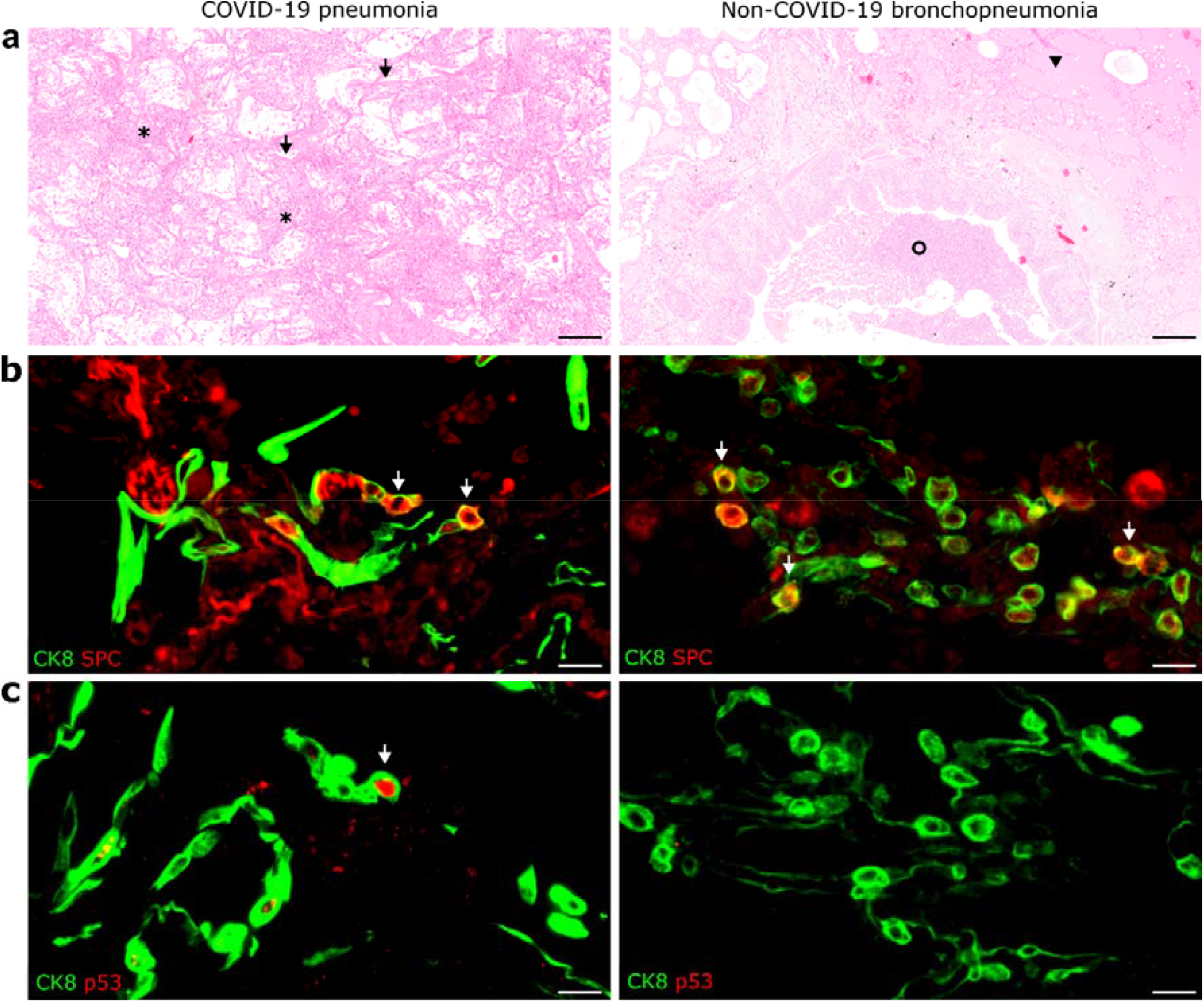
Alveolar differentiation intermediate (ADI) cells in COVID-19 and non-COVID-19 pneumonia. A Representative images showing histopathological lesions in a COVID-19 patient (left) and in non-COVID-19 bronchopneumonia case (right). COVID-19 is characterized by diffuse alveolar damage (DAD) with hyaline membranes (arrows) and alveolar spaces filled with sloughed epithelial cells, leukocytes and edema (asterisks). Non-COVID-19 bronchopneumonia was characterized by intraluminal suppurative exudate (circle) and alveolar edema (arrowhead) without DAD. **B** Representative image of double immunofluorescence for the ADI marker CK8 (green) and the AT2 marker proSP-C (red) in a COVID-19 (left) and non-COVID-19 bronchopneumonia (right) sample. Cells with a round morphology express both markers (arrows). **C** Representative image of double immunofluorescence for the ADI marker CK8 (green) and the cell cycle exit marker TP53 (red) in a COVID-19 (left) and a non-COVID-19 bronchopneumonia (right) sample. ADI cells in COVID-19 patients express TP53(arrow), while ADI cells in the non-COVID-19 bronchopneumonia case are negative. Scale bars: 200 µm and 20 µm (b, c).

In conclusion, ADI cells are a feature of alveolar regeneration following SARS-CoV-2 induced DAD in lethal COVID-19 and its Syrian golden hamster model. These cells were also detected in low numbers under non-infectious conditions in the hamster and in a human sample with suppurative bronchopneumonia, confirming that ADI cells participate in physiological turnover and alveolar repair regardless of the etiology in both species. Importantly, only ADI cells from SARS-CoV-2 infected hamsters and humans expressed TP53, hinting at a prolonged block of these cells in the intermediate state.

### 3. Multipotent airway-derived CK14^+^ progenitors contribute to alveolar regeneration following SARS-CoV-2 induced DAD in hamsters

It is well accepted that upon severe alveolar injury, both AT1 and AT2 cells can be replenished by airway progenitors (**Fig. 4 A)** ^15,17,38-40^. In the next step, we characterized the contribution of airway progenitors to alveolar regeneration in SARS-CoV-2 infected hamsters. As described earlier, histopathological lesions at 6 and 14 dpi included foci of prominent alveolar epithelial proliferation with airway-like morphology that were frequently in anatomic continuity with bronchiolar-alveolar junctions. Thus, we determined 1) the cellular origin of these proliferates and 2) whether these progenitors differentiate into AT2 or ADI cells after migrating into the alveoli.

**Figure 4:**
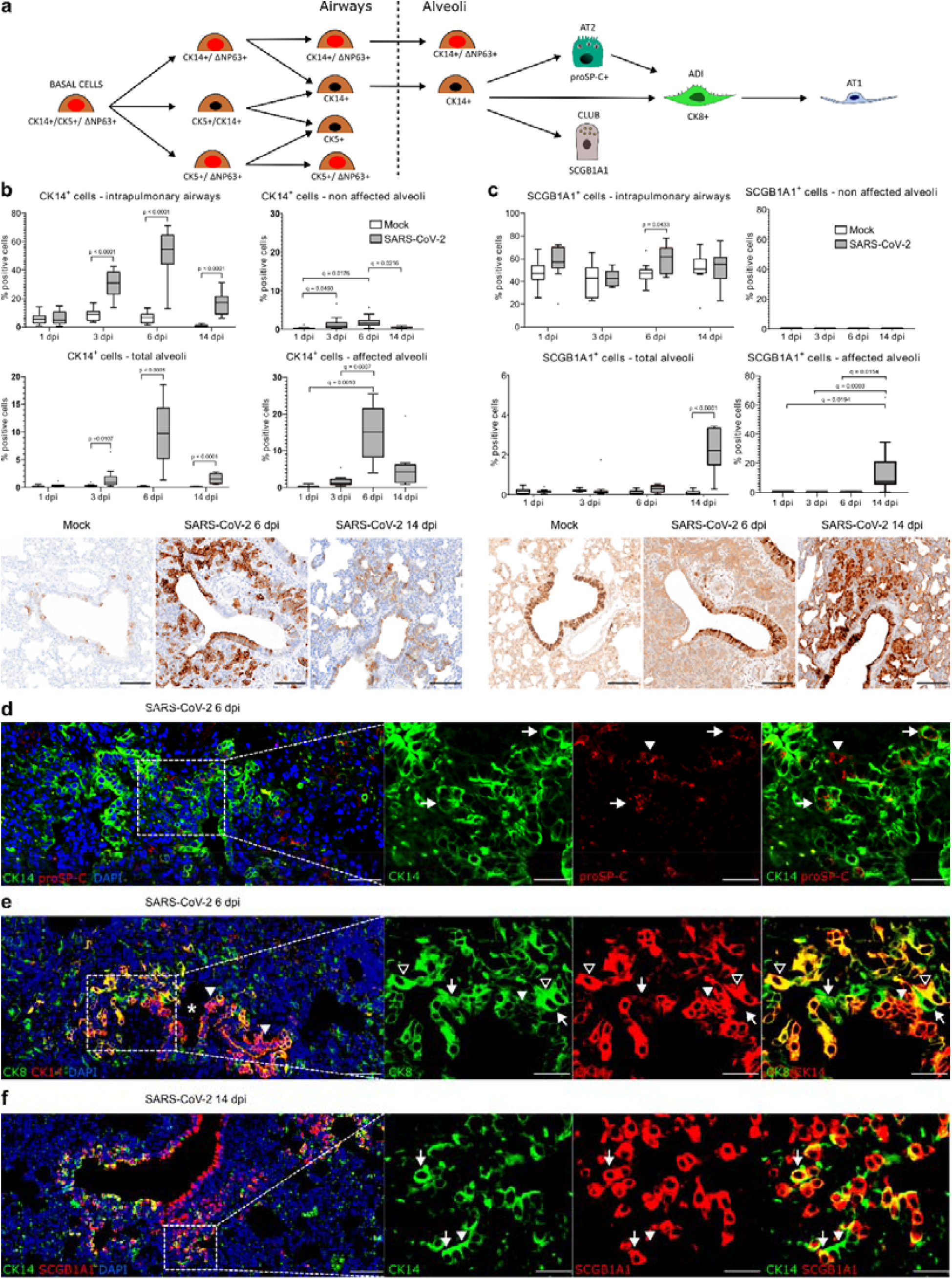
Airway basal cells participate in alveolar regeneration in SARS-CoV-2 infected hamsters. A Schematic illustration of the proposed trajectory of airway basal cells towards alveolar cells. CK14^+^CK5^+^ΔNP63^+^ basal cells proliferate within the airways and give rise to different combinations of CK5^+/-^, CK14^+/-^, ΔNp63^+/-^ progenitor cells (see also supplementary Fig. 5). Upon severe alveolar damage, rare ΔNp63^+^CK14^+^ and frequent CK14^+^ basal cells mobilize to the alveoli giving rise to alveolar pneumocytes type 2 (AT2, see D) and/or to alveolar differentiation intermediate (ADI) cells (see E), particularly at 6 dpi. At 14 dpi, CK14^+^ basal cells give rise to secretoglobin 1A1^+^ (SCGB1A1) club cells within the peribronchiolar alveolar proliferates (see F). **B, C** Quantification of CK14^+^ basal cells (B) and SCGB1A1^+^ club cells (C) within intrapulmonary airways, total alveoli, non-affected alveoli, and affected alveoli as well as representative pictures of immunolabeled cells (brown signal) in the bronchioles and peribronchiolar proliferates in mock and SARS-CoV-2 infected hamsters at 6 and 14 dpi. The percentage of the immunolabelled cells relative to total cells in the respective area is given. Pictures of SARS-CoV-2 infected hamsters at 6 and 14 dpi are taken from the same location for CK14 and SCGB1A1 immunolabelings. **D** Representative image of double immunofluorescence for CK14 (green) and proSP-C (red) in a peribronchiolar proliferation area in a SARS-CoV-2 infected hamster at 6 dpi. An overview and higher magnification of the area delineated by a rectangle are shown. The arrowhead shows a proSP-C^+^ AT2 cell. The arrows indicate double labeled airway progenitors differentiating into proSP-C^+^ AT2 cells. **E** Representative image of double immunofluorescence for CK14 (red) and CK8 (green) in a peribronchiolar proliferation area in a SARS-CoV-2 infected hamster at 6 dpi. An overview and higher magnification of the area delineated by a rectangle are shown. The image shows a transition from CK14^+^ airway basal cells forming a pod (white arrowhead), to double labeled CK14^+^CK8^+^ cells differentiating into elongated ADI cells (open arrowheads) and CK14^-^CK8^+^, elongated ADI cells (arrows). **F** Representative image of double immunofluorescence for CK14 (green) and SCGB1A1 (red) in a peribronchiolar proliferation area in a SARS-CoV-2 infected hamster at 14 dpi. An overview and higher magnification of the area delineated by a rectangle are shown. A transition from CK14^+^ airway basal cells (arrowhead) to CK14^+^SCGB1A1^+^ club cells (arrows) is shown. Quantification data are shown as box and whisker plots. Statistical analysis was performed by two-tailed Mann-Whitney-U test. For multiple comparisons between time points, a Benjamini–Hochberg correction was applied. P- and q-values ≤0.05 were considered significant. ND=D10 animals/group for mock and SARS-CoV-2 respectively. For quantifications, 1 longitudinal section containing all right lung lobes were evaluated. Source data will be provided as a source data file. Scale bars: 100 µm (c, d), 50 µm (overview in d, e, f), 25 µm (high magnification in d, e, f).

Multiple airway progenitor cell types have been reported to contribute to alveolar regeneration, including proSP-C^+^SCGB1A1^+^ broncho-alveolar stem cells (BASCs), ΔNP63^+^CK5^+^ distal alveolar stem cells (DASCs), ΔNP63^+^CK5^+^CK14^+^ basal cells, and SCGB1A1^+^ club cells ^40-42^. First, our aim was to identify these cell types in the distal airways of hamsters. The predominant basal cell type was CK14^+^, followed by CK14^+^ΔNP63^+^ cells (Suppl fig 5). ΔNP63^+^CK5^+^CK14^+^ cells were rare in the distal airways (Suppl fig 5). We did not detect ΔNP63^+^CK5^+^ DASCs, CK5^+^ cells or SPC^+^SCGB1A1^+^ BASCs in the distal airways of hamsters (*data not shown*). In addition to basal cell types, SCGB1A1^+^ club cells were detected in high numbers in distal airways.

In the peri-bronchiolar proliferation foci of SARS-CoV-2 infected animals at 6 dpi, the majority of cells were CK14^+^, while CK14^+^ΔNP63^+^ cells were rare (Supplem. Fig. 6). CK5^+^, CK5^+^ΔNP63^+^ or CK14^+^CK5^+^ cells were not detected within these areas (Supplem. Fig. 6). SCGB1A1 expression was absent in the peri-bronchiolar proliferates at 6 dpi, but abundantly present at 14 dpi. Therefore, we focused our further quantitative analysis on CK14^+^ airway basal cells and SCGB1A1^+^club cells.

In mock-infected hamsters, the number of CK14^+^ cells in the airways remained unchanged over the observation period (**Fig. 4 B)**. SARS-CoV-2 infection caused a marked proliferation of CK14^+^ cells in the airways, which peaked at 6 dpi and remained elevated until 14 dpi. The number of CK14^+^ cells in total alveoli was significantly increased compared to the mock group at 3, 6 and 14 dpi, mirroring the increase in the airways (**Fig. 4 B)**. CK14 was expressed by the majority of cells in the peri-bronchiolar proliferation forming pods and tubules continuous with terminal bronchioles at 6 dpi. At 14 dpi, the peri-bronchiolar proliferates were only partly CK14^+^ (**Fig. 4 B)**.

In contrast to the CK14^+^ progenitors, we observed no major contribution of club cells in the alveolar proliferative response during early infection (**Fig. 4 C)**. The number of SCGBA1^+^ club cells in the airways remained similar in mock-infected animals at all time-points. In SARS-CoV-2 infected animals, the number of SCGB1A1^+^ cells in the airways was mildly increased compared to mock at 6 dpi (**Fig. 4 C)**. SCGB1A1 was not expressed in alveolar proliferation areas at 3 and 6 dpi. Interestingly, SCGB1A1^+^ cells significantly increased in the alveoli of SARS-CoV-2 infected animals at 14 dpi. The expression was limited to the airway-like, peri-bronchiolar proliferation areas, in which up to 40% of cells were SCGB1A1^+^ club cells (**Fig. 4 C)**.

Therefore, it was concluded that CK14^+^ cells are the airway progenitors that mainly contribute to alveolar regeneration in SARS-CoV-2 infected hamsters. These cells probably have their origin in a common ΔNP63^+^CK5^+^CK14^+^ basal cell pool, but represent a subset that loses CK5 and partly ΔNP63 expression upon migration into the alveoli.

Next, we determined the fate of the CK14^+^ cells in the alveoli. Double-labeling with proSP-C revealed clusters of CK14^+^proSP-C^+^ cells in the peri-bronchiolar pods and occasionally within the lining of terminal bronchioles. This indicates a potential differentiation of airway progenitors towards the AT2 lineage (**Fig. 4 D;** Supplem. Fig. 7 A-B). At the edges of the peri-bronchiolar proliferates, some CK14^+^ cells showed a transition from a cuboidal to an elongated shape typical of ADI cells. Co-staining with CK8 showed a gradual phenotypical change in the direction of alveoli. Cells exiting the bronchiole showed a cuboidal morphology and a diffuse cytoplasmic CK14 expression. Towards alveoli, the cuboidal cells co-expressed CK14 and CK8. More distally, cells became more elongated and were characterized by CK14^-^CK8^+^ immunolabeling (**Fig. 4 E;** Supplem. Fig. 7 C-D). Therefore, we concluded that airway progenitors can differentiate into AT2 but also directly into the ADI state. These transitions were mainly observed at 6 dpi. In contrast, at 14 dpi, peri-bronchiolar CK14^+^ cells partly co-expressed SCGB1A1, indicating a club cell differentiation (**Fig. 4 F;** Supplem. Fig. 7 E-F). Hence, we concluded that the increased number of alveolar SCGB1A1^+^ cells we observed at this time point was most likely the result of *in situ* differentiation of CK14^+^ cells. However, we cannot exclude that SCGB1A1^+^ club cells also proliferated and migrated out of the bronchioles to give rise to alveolar cells at 14 dpi.

In summary, our findings indicate that multipotent CK14^+^ airway basal cell progenitors, probably arising from a CK14^+^CK5^+^ΔNP63^+^ basal cell pool, proliferate and migrate to alveoli following SARS-CoV-2 induced DAD in hamsters. These cells have the potential to differentiate into distinct lineages, including AT2, ADI and club cells, depending on the timing and localization.

### 4. Hamsters show dysregulated alveolar regeneration and fibrosis following SARS-CoV-2 induced DAD

SARS-CoV-2 NP antigen was no longer detectable in the lung at 6 dpi. However, ADI cells and airway progenitors were still present in the alveoli at 14 dpi, indicating an ongoing regeneration processes with incomplete restoration of alveolar structures at this time-point. Moreover, 7 out of 9 animals showed multifocal, sub-pleural, variably sized, well demarcated areas with aggregates of spindle cells and abundant, pale, fibrillary, extracellular material (**Fig. 5 A)**. Azan staining confirmed deposition of collagen in these areas (**Fig. 5 B)**. Immunohistochemistry for α-smooth muscle actin (α-SMA) demonstrated the presence of myofibroblasts (**Fig. 5 C)**. The fibrotic areas encompassed from 0.59 to 2.35 % of the evaluated lung tissue area (**Fig. 5 E)**. Next, we wanted to determine if ADI cells and fibrosis at 14 dpi are locally associated with M2-polarized macrophages in hamsters. CD204^+^ M2-type macrophages were frequently detected within and around fibrotic areas (**Fig. 5 D)**. The number of CD204^+^ cells was significantly higher in SARS-CoV-2 infected animals compared to the mock group at 3, 6 and 14 dpi (**Fig. 5 F)**.

**Figure 5:**
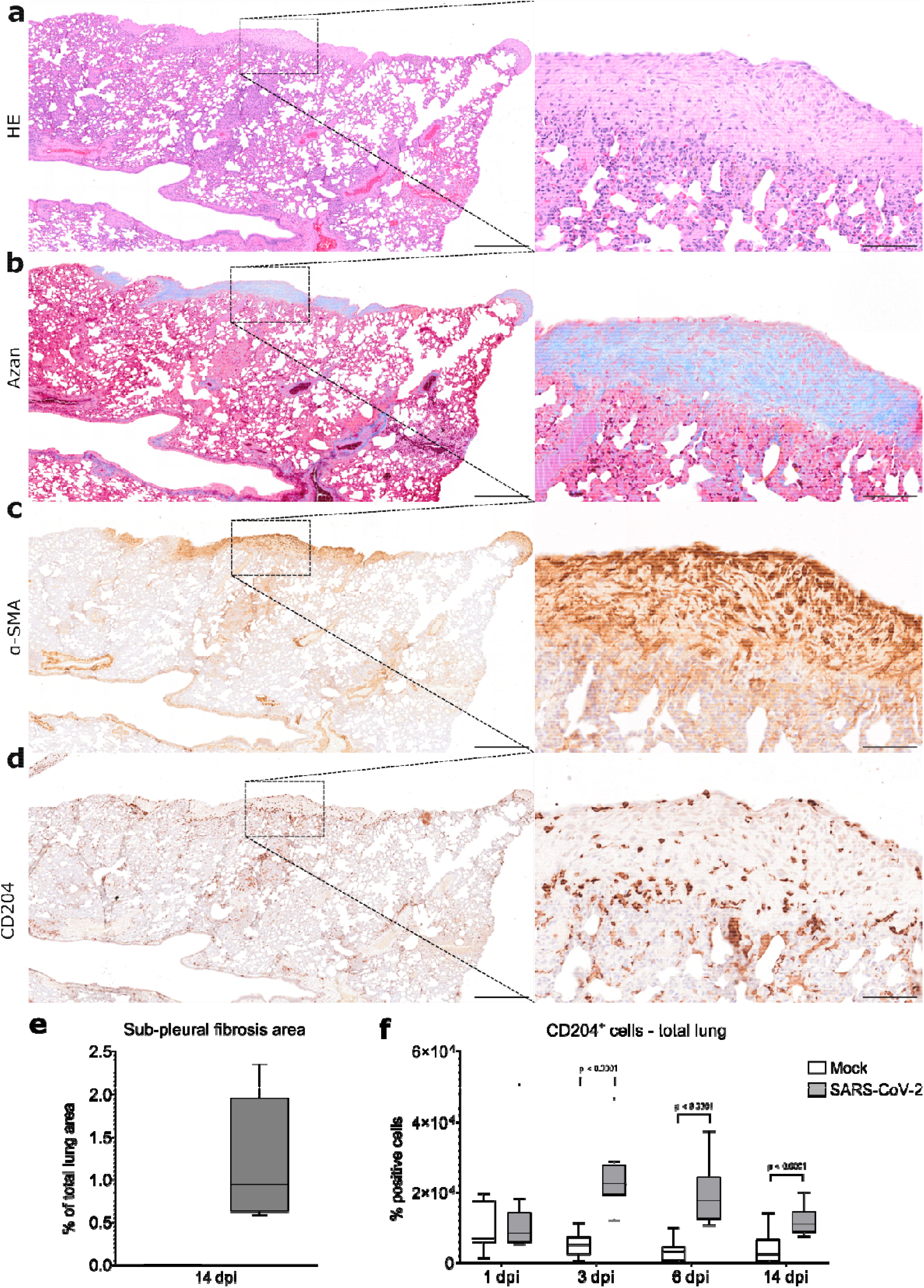
Sub-pleural fibrosis in SARS-CoV-2 infected hamsters. **A-D** Representative images showing sub-pleural fibrotic foci in a lung lobe of a SARS-CoV-2 infected hamster at 14 dpi. The left panel shows an overview of one right lung lobe displaying multifocal, extensive, well demarcated areas of sub-pleural fibrosis. The right panel shows at higher magnification of the area delineated by the rectangle. On hematoxylin-eosin (HE) stained sections, this lesion is characterized by sub-pleural aggregates of spindle cells and abundant, pale eosinophilic, fibrillary, extracellular matrix (A). Azan stain demonstrates the presence of mature collagen fibers in the matrix (blue signal, B). Immunohistochemistry shows abundant α-smooth muscle actin (α-SMA)^+^ myofibroblasts (brown signal in C) as well as infiltration with CD204^+^ M2 macrophages (brown signal, D). **E** Quantification of sub-pleural fibrosis in lungs of mock and SARS-CoV-2infected hamsters at 14 dpi. The percentage of affected area relative to total lung area is given. **F** Quantification of CD204^+^ M2 macrophages in total lung area. Data are shown as box and whisker plots. Data from CD204 quantification was tested by two-tailed Mann-Whitney-U test. A p-value of ≤0.05 was chosen as the cut-off for statistical significance. N□=□10 animals/group for mock and SARS-CoV-2 respectively. For quantifications, 1 longitudinal section containing all right lung lobes were evaluated. Source data will be provided as a source data file. Scale bars (a-d): 500 µm (overview in a-d), 100 µm (high magnification in a-d).

In summary, the findings revealed an incomplete restoration of alveolar structures with ADI cells and M2-type macrophages, as well as sub-pleural fibrosis, still detectable two weeks after infection.

### 5. Single-cell transcriptome analysis confirms ADI cell persistence following SARS-CoV-2 induced DAD in hamsters

As described above, we demonstrated that ADI cells with features previously described in mouse models of lung regeneration as well as in COVID-19 patients are participating in alveolar regeneration following SARS-CoV-2 infection of hamsters. To confirm this observation with data from an independent experiment, we re-analyzed a previously published single-cell RNASeq dataset (GSE162208) generated in SARS-CoV-2 infected Syrian golden hamsters ^43^. The experiment was performed with a study design similar to the present investigation. We focused our analysis on data from SARS-CoV-2-infected animals sacrificed at 5 and 14 dpi. First, we generated a Uniform Manifold Approximation and Projection (UMAP) clustering all cell populations detected in the datasets. We then identified alveolar cells based on the expression of AT1 and AT2 markers (*Rtkn2* and *Lamp3*, respectively), as described in the original publication (**Fig. 6 A, G**)^43^. These cells were re-clustered according to differences in gene expression, resulting in 7 and 11 clusters at 5 and 14 dpi, respectively. Next, we determined the top 10 differentially expressed genes (DEGs) in each cluster and compared the sets of DEGs with gene signatures described in mouse models of lung regeneration ^19,20^ as well as COVID-19 patients ^9^. Within the DEGs, we detected genes typically expressed by AT1, AT2, ADI cells, club cells or ciliated cells in mice and/or humans, and we generated lists of candidate marker genes for these cell types in the hamster. Next, we evaluated the expression of these candidate markers within the clusters and removed genes with low specificity from the lists. The final, hamster-specific marker gene lists are given in supplementary table 1. The module scores of the respective marker sets at 5 and 14 dpi are visualized in **Fig. 6 B-F; H-L**.

**Figure 6:**
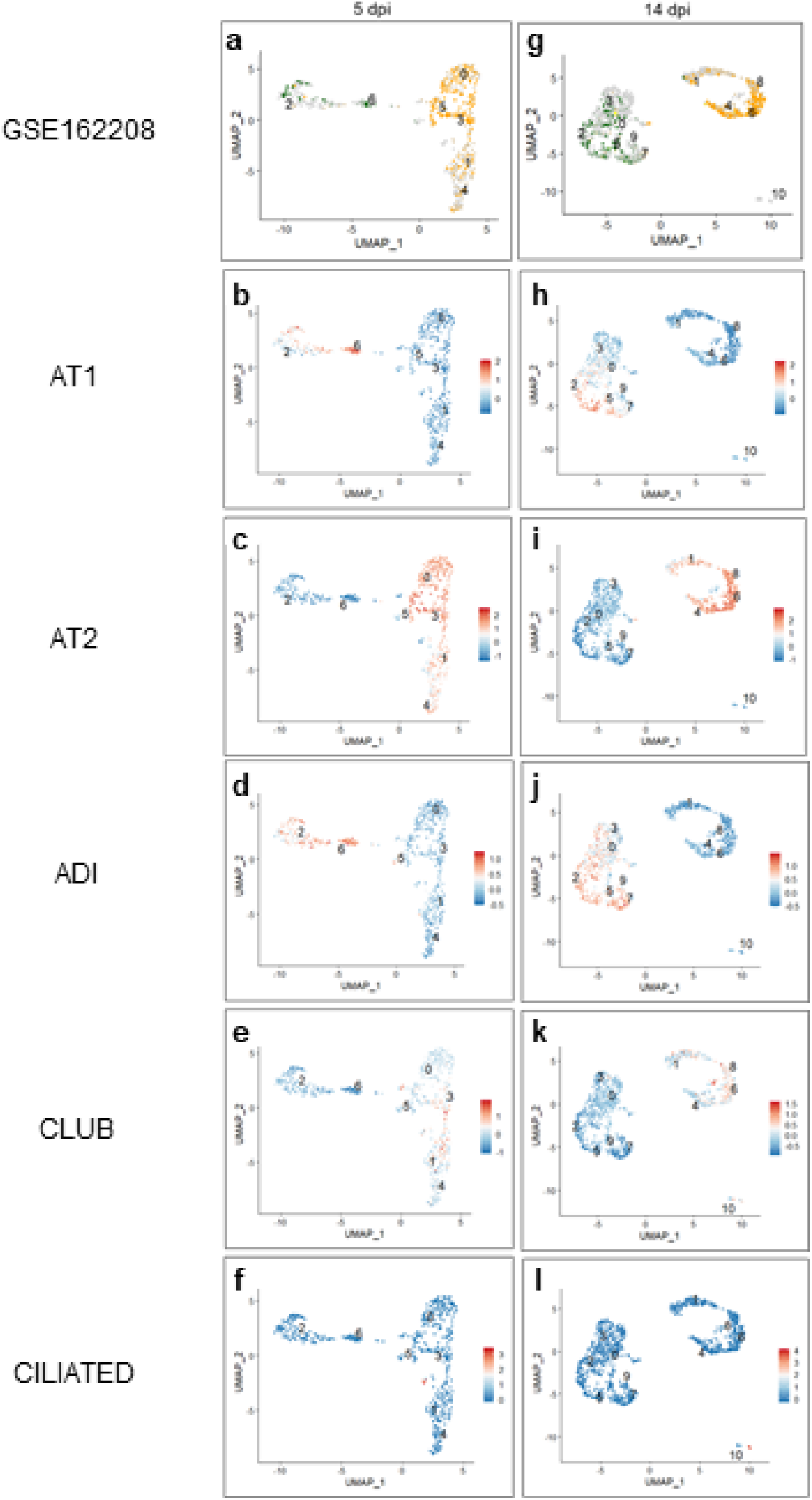
Single cell analysis of alveolar cells in SARS-CoV-2 infected hamsters. Single cell RNA-Seq data set (GSE162208) from lungs of SARS-CoV-2 infected hamsters killed at 5 (A-F) or 14 (G-L) days post infection (dpi). **A, G** Expression of AT1 (green *Rtkn2*) and AT2 (orange *Lamp3*) marker genes. **B-F** and **H-L** Results from module score analysis for cell marker genes. For cell marker gene list, see supplementary table 1.

At 5 dpi, the AT1 marker *Rtkn2* was expressed in a small number of cells in clusters 2 and 6 (**Fig. 6 A**). The AT2 marker *Lamp3* was mostly expressed in many cells within a separate cell population, comprised of clusters 0, 1, 3, 4 and 5. Interestingly, *Lamp3* was also detected in some cells within cluster 2, indicating a mixed composition of this cluster (**Fig. 6 A**). Many cells did not express any one of the two genes. Applying the module scores algorithm with sets of multiple marker genes allowed a distinction of mature and transitional alveolar cell types. Mature AT1 marker genes scored high in cluster 6 and partly in cluster 2, in line with the distribution of *Rtkn2* expression (**Fig. 6 B**). Mature AT2 genes showed positive scores in clusters 0, 1, 3, 4 and 5 (**Fig. 6 C**), but not within the AT1 clusters. Positive scores for ADI marker genes were detected throughout clusters 2 and 6 and partly in cluster 5 (**Fig. 6 D**). Importantly, high scores were observed in the cells that did not score for markers of mature AT1 and AT2 cells. Interestingly, clusters showing high expression of AT2 genes also partly showed high scores for club cell genes (cluster 1, and 3, **Fig. 6 E**). A group of cells within cluster 5 only scored high for club cell genes (**Fig 6 E**). A few cells within cluster 1 scored high exclusively for gene markers of ciliated cells (**Fig. 6 F**).

At 14 dpi, the AT1 marker *Rtkn2* was expressed in clusters 0, 2, 3, 5, 7 and 9. The number of *Rtkn2*-positive cells was higher compared to 5 dpi. Similar to 5 dpi, *Lamp3* was expressed in a separate population (clusters 1, 4, 6, and 8) and also partly within the AT1 cell clusters (**Fig. 6 G**). Again, many cells were negative for both genes. Module scores for AT1 genes were high only in three clusters expressing *Rtkn2* (2, 5 and 7, **Fig. 6 H**). AT2 genes scored high in 4 clusters (1, 4, 6 and 8, **Fig 6 I**). Interestingly, the majority of cells within three clusters (0, 3 and 9) showed no positive scores for either AT1 or AT2 gene sets, but scored high for ADI marker genes (**Fig 6 J**). The number of cells with high scores for ADI cell genes was higher compared to 5 dpi. Similar to 5 dpi, a positive score for club cell genes was detected within AT2 clusters (cluster 6 and 8, **Fig. 6 K**). In addition, a positive score for club cells genes was observed in some cells within one of the ADI cell clusters (cluster 3). Cluster 10 separated completely from the other populations and showed a high score for ciliated cell markers (**Fig. 6 L**).

Taken together, transcriptome analysis identified AT1, AT2 and ADI cells in SARS-CoV-2-infected hamsters. At 5 dpi, ADI cells did not form a separate cluster, but were admixed with AT1 and AT2 cells. At 14 dpi, ADI cells were more numerous and clustered separately from AT1 and AT2 cells. Moreover, we found small groups of ciliated cells admixed within the alveolar cell populations and partial expression of club cell genes within ADI and AT2 cells.

Next, we wanted to investigate the expression of genes belonging to pathways involved in lung regeneration and we performed module score analysis with hallmark gene lists (http://www.gsea-msigdb.org/gsea/msigdb/index.jsp): *p53 pathway, DNA repair, TGF beta signaling, notch signaling, wnt beta catenin signaling, epithelial mesenchymal transition (EMT), and angiogenesis*. As described above, ADI cells in mice and humans express Tp53 and other markers of cell cycle arrest and DNA repair. The transcriptome data showed that a fraction of cells with an ADI signature showed high positive scores for *p53 pathway* genes at 5 and 14 dpi (**Fig 7 A,B**). At 5 dpi, almost all clusters showed positive scores for *DNA repair* genes, with the highest scores observed in AT1/ADI and cells with a ciliated cell signature (**Fig 7 C**). At 14 dpi, mainly AT1/ADI and ADI clusters displayed positive scores (**Fig. 7 D**). The AT2-ADI-AT1 trajectory is regulated by different signaling pathways, including TGF beta -, notch - and wnt beta catenin signaling and involves the EMT process ^19,44^. At 5 and 14 dpi, only a few cells with ADI signature showed high positive scores for *TGF beta signaling* (**Fig. 7 E-F**). A minimal number of AT2 cells revealed a high positive score for *notch signaling* hallmark genes at 5 dpi, whereas variably positive scores were distributed within AT2, ADI and AT1/ADI cells at 14 dpi (**Fig. 7 G-H**). A small number of cells within the AT1/ADI cluster revealed high positive scores for *wnt beta catenin signaling* hallmark genes at 5 dpi (**Fig. 7 I**). At 14 dpi, larger numbers of cells within AT1/ADI and ADI clusters as well as a small number of cells within the AT2 clusters showed positive scores for *wnt beta catenin signaling* hallmark genes (**Fig. 7 J**). AT1/ADI clusters as well as cells with an ADI signature within the AT2 clusters showed a high positive score for *EMT* hallmark genes at 5 dpi (**Fig. 7 K**). At 14 dpi, some AT1/ADI cells showed positive scores for *EMT* hallmark genes (**Fig. 7 L**). Finally, we investigated the expression of genes involved in angiogenesis, since this process is upregulated in late phases of DAD, in the context of fibrosis ^45^. A small number of cells within the AT1/ADI cluster at 5 dpi and a higher number of cells within the AT1/ADI and ADI clusters at 14 dpi revealed high positive scores for *angiogenesis* hallmark genes (**Fig. 7 M-N**).

**Figure 7:**
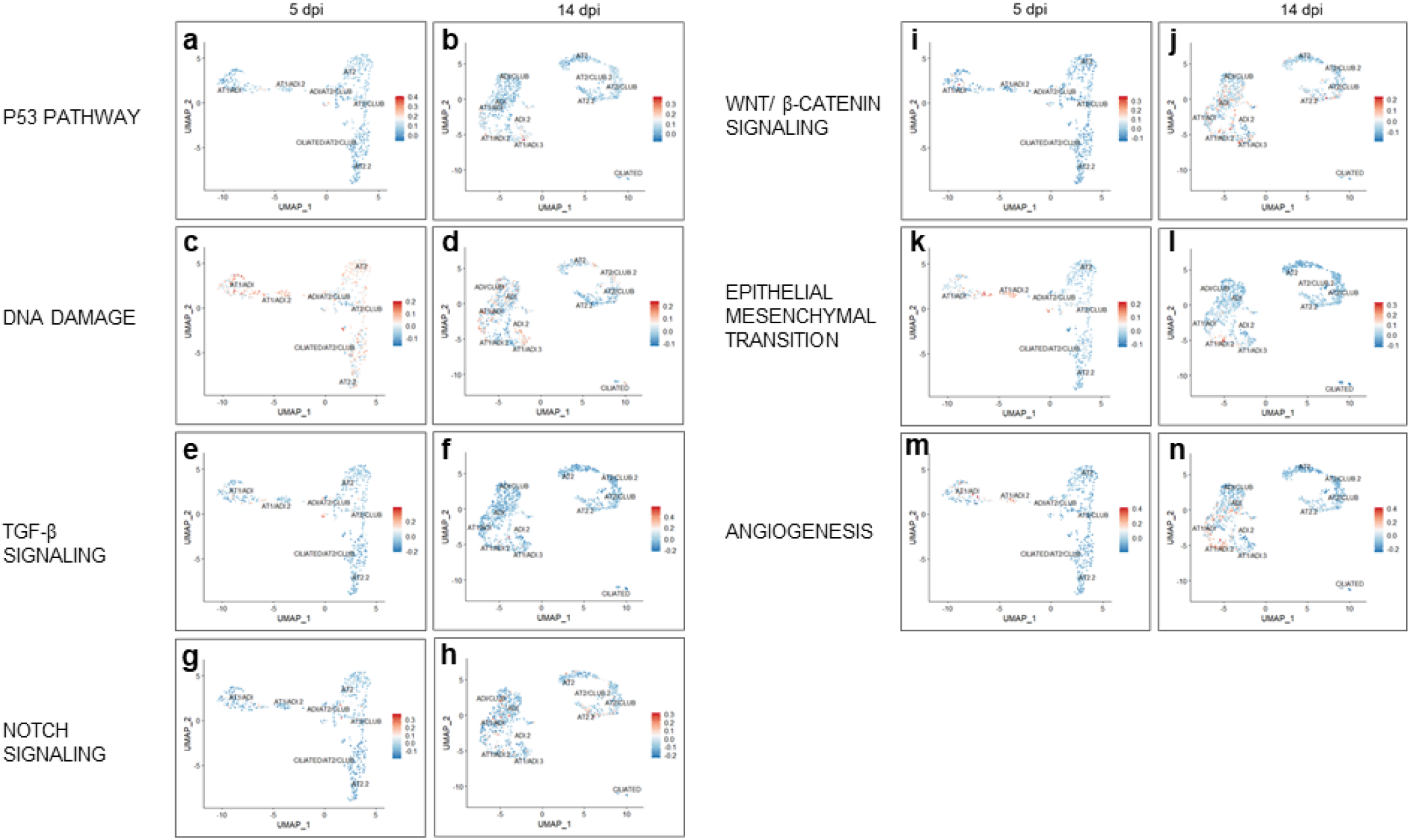
Module scores for GSEA hallmark genes within alveolar cells in SARS-CoV-2 infected hamsters. **A-N** Results from module score analysis for GSEA hallmark genes. The cluster names are indicated.

In summary, the findings from the independent study confirmed that ADI cells are a feature of alveolar regeneration in hamsters on a transcriptome level, supporting the morphologic observations from our experiment. Moreover, the data shows that i) the number of AT1 cells increased from 5 to 14 dpi, indicative of progressive alveolar regeneration, ii) cells with an ADI gene signature can be distinguished within AT1 and AT2 populations and they become more distinct and numerous at 14 dpi, iii) ADI cells partly express genes belonging to the *p53* and *DNA repair pathway* as well as *TGF beta* -, *notch*- and *wnt beta catenin* signaling, *EMT and angiogenesis pathways* iv) club cell genes are partly expressed in AT2 and ADI cells at 14 dpi in SARS-CoV-2 infected hamsters.

## Discussion

The COVID-19 pandemic has claimed many lives and challenged the global healthcare system in an unprecedented way. Survivors of acute disease may be faced with a wide spectrum of long-lasting symptoms, with pulmonary, neuropsychiatric and cardiovascular sequelae at the forefront, which have a negative impact on the quality of life. Considering the staggering amount of patients reporting prolonged symptoms even as long as 15 months after the initial onset of COVID-19 ^2,7,46,47^, further research into potential pathomechanisms of this protracted recovery is urgently needed ^48^. A possible explanation for the mechanisms underlying some PASC symptoms, such as dyspnea, shortness of breath and exercise intolerance, could be an impaired regeneration of alveolar tissue and lung fibrosis ^3,9^. It has also been suggested that the persistence of CK8^+^ ADI cells might be the cause of prolonged hypoxemia in COVID-19 patients ^8^. Importantly, these conclusions are based on observations from tissues collected from acute, lethal COVID-19 cases. In contrast, we can only speculate about the presence of these cells in PASC, since samples from affected humans are scarce. Therefore, establishment and further characterization of appropriate animal models of PASC are urgently needed.

SARS-CoV-2 infected hamsters reliably phenocopy moderate to severe COVID-19 ^28^. Recovering hamsters show a pronounced epithelial cell proliferation within airways and alveoli, which started at 3 dpi and was detectable until 14 dpi in the present study. This is in line with previous reports, which showed that proliferative foci can persist up to 31 dpi in hamsters^33,49^. Here, we characterized the proliferating epithelial cell types in more detail. First, we investigated the AT2-ADI-AT1 trajectory. In mouse models of lung injury, the transition from AT2 to the ADI state is characterized by progressive decrease of cell sphericity, expression of CK8 and loss of AT2 marker expression ^8,19-21^. Double-labelling of SPC and CK8 demonstrated the transition of AT2 to ADI cells, associated with phenotypical changes as described above in SARS-CoV-2-infected hamsters. At 6 dpi, all stages of ADI cells were observed, including round, SPC^+^CK8^+^ cells (early ADI stage) and polygonal, plump to elongated, SPC^-^CK8+ cells (late ADI stage). Interestingly, at 14 dpi, we observed numerous round, SPC^+^CK8^+^, early ADI stages and fewer late ADI stages, which could indicate a new wave of ADI cell generation at this time-point. Lineage tracing studies in the mouse bleomycin lung injury model demonstrated that ADI cells could develop from AT2 as well as from MHCII^+^ club cells migrating from the airways ^19^. In early stages after injury, peaking at 5 dpi, ADI cells are mainly derived from AT2 cells, while club cell-derived ADI cells appear later, peaking at 10 dpi. Of note, a part of the MHCII^+^ club cells differentiating towards ADI cells goes through an SPC^+^ stage ^19^. We speculate, that the round SPC^+^CK8^+^ ADI cells observed at 14 dpi in SARS-CoV-2 infected hamsters could be derived from airway progenitors analogous to murine MHCII^+^ club cells, which transiently assume an AT2 stage.

In addition to the demonstration of transitional cell stages on a morphological level, the presence of ADI cells in the hamster model of COVID-19 was confirmed using transcriptome data analysis. In addition, we created hamster-specific marker gene lists for different alveolar cell populations, including AT1, ADI and AT2 cells. Importantly, numerous cells with an ADI gene signature were detected at 14 dpi, which is indicative of an ongoing regenerative process at this time point and in line with the results obtained by the quantification of CK8 positive cells by immunolabeling. At 5 dpi, cells with ADI gene expression clustered with AT1 and AT2 cells, suggestive of an AT2 origin. Interestingly, at 14 dpi, ADI gene expression was not found within the AT2 clusters. At this time point, a small number of ADI cells expressed a club cell signature. This observation reinforces the hypothesis, that two waves of ADI cells are generated in the course of SARS-CoV-2 infection of hamsters, which have their origin in AT2- and club cells, respectively.

CK8^+^ADI cells in SARS-CoV-2 infected hamsters frequently expressed nuclear TP53 protein. Transcriptome data also showed that some cells with ADI gene signature displayed high scores for *p53 pathway* and for *DNA repair* hallmark genes at 5 and 14 dpi. Nuclear TP53 regulates transcription of genes involved in cell cycle arrest and DNA repair and accumulation of TP53 is therefore detected in cells with high level of DNA damage ^50^. ADI cells undergo mechanical stretch-induced DNA damage while migrating to cover the denuded septa and to differentiate into AT1 ^21,51^. The nuclear expression of TP53 could reflect a particularly high level of injury, triggering DNA repair mechanisms. It is important to underline that in SARS-CoV-2 infected hamsters, nuclear TP53 expression was often found in hypertrophic CK8^+^ cells with a bizarre morphology, binucleation or karyomegaly. We assume that these hypertrophic cells have accumulated a high level of DNA damage, are blocked in the ADI stage and are not likely to differentiate into slender AT1 cells. A permanent block in the ADI cell state has been described in idiopathic pulmonary fibrosis (IPF) and mouse models of lung fibrosis ^8,20,52,53^. Importantly, it has been demonstrated in a mouse model that induction of TP53-dependent AT2 senescence is sufficient to propagate progressive pulmonary fibrosis ^44,53^. Besides TP53, other signaling pathways have been implicated in ADI cell senescence. For instance, *in vitro* studies in primary murine cells revealed that a chronic activation of WNT/β-catenin signaling can induce senescence and CK8 expression in ADI cells ^44,54^. In addition, persistent Notch activation in AT2 cells induces retarded differentiation of AT2 into AT1 cells, resulting in ADI cell accumulation in a *Pseudomonas* lung injury model ^44,55^. Moreover, persistent TGF-β signaling has been shown to block ADI cells from differentiating into AT1 cells ^20^. We showed that, from 5 to 14 dpi, an increasing number of cells with ADI gene signature had high scores for *Wnt/* β*-catenin* and *notch signaling* hallmark genes. In contrast, genes belonging to the TGF-β signaling pathway showed no high scores at 14 dpi and were only detected in a small fraction of ADI cells at 5 dpi. Therefore, we speculate that prolonged Wnt/ β-catenin and/or notch signaling, rather than excessive TGF-β, could be responsible for the prolonged presence of ADI cells in SARS-CoV-2 infected hamsters. However, the available data do not allow us to assess the duration of the activation of the respective pathways in ADI cells and further studies with a more detailed analysis and additional time points are warranted to confirm this hypothesis. Besides dysregulation of the discussed pathways, a direct contribution of viral infection to the induction of senescence must be considered. It has been demonstrated that SARS-CoV-2 and other viruses can induce cellular senescence in infected AT2 cells ^56^.

The clinical relevance of the observed ADI cell accumulation in hamsters deserves further investigation. In COVID-19 patients with a severe disease course and lethal outcome, high numbers of ADI cells were detected by others and in the present study, which indicates that dysregulated alveolar regeneration could play a role in the pathogenesis of severe disease ^8,9^. In line with this, we found that TP53 is expressed by CK8^+^ ADI cells in lethal COVID-19 samples, but not in CK8^+^ ADI cells in a non-COVID pneumonia case.

In addition to the presence of ADI cells, the majority of SARS-CoV-2 infected animals showed foci of sub-pleural fibrosis at 14 dpi, indicative of irreversible damage/remodeling. This is in line with previous reports in hamsters ^57,58^. The pattern of fibrosis is similar to what has been described in IPF patients and a RhoGTPase Cdc42 deletion mouse model of progressive pulmonary fibrosis ^23,59^. In these conditions, a progression of fibrotic lesions from periphery to center is typically encountered ^19,23^. Subpleural alveoli are subject to increased mechanical tension during respiration, which has been shown to activate TGF-β-mediated pro-fibrotic processes ^23,59^. In addition to fibrotic foci, our study also revealed a prominent presence of CD204^+^ M2 macrophages starting at 3 dpi and persisting until 14 dpi. M2-macrophages are known to promote fibrosis by a variety of factors, including TGF-β secretion ^60^. Thus, the fibrosis could be promoted by the prolonged presence of an unfavorably polarized inflammatory response. In addition to macrophages, AT2 cells can promote a pro-fibrotic microenvironment by activating local fibroblasts to become myofibroblasts via paracrine signaling, as demonstrated *in vitro* ^45,61,62^. This process was initiated by an induction of an EMT process in the AT2 cells ^53,62^. Of note, it has been reported that EMT is activated in ADI cells ^19^ and the results from our transcriptome analysis showed that cells with ADI gene signature score high for EMT pathway gene expression at 5 and 14 dpi in SARS-CoV-2 infected hamsters. Therefore, besides M2-macrophages, ADI cells potentially contribute to a pro-fibrotic microenvironment. In addition, it has been reported that lung fibrotic lesions in COVID-19 patients are preceded by a prolonged blood vessel neo-formation ^63^. Interestingly, transcriptome data revealed that numerous cells within AT1/ADI and ADI cell clusters showed high positive scores for *angiogenesis* hallmark genes at 14 dpi, suggesting that ADI cells in hamsters might contribute also to a pro-angiogenetic microenvironment, promoting vascular changes during lung fibrosis similar to COVID-19 patients. A recent study in a mouse model of COVID-19 demonstrated that aged mice infected with a mouse adapted strain of SARS-CoV-2 show fibrotic lesions starting from 15 dpi and persisting up to 120 dpi ^52^. Similar to what we observed in the hamster model, the lesions were characterized by a subpleural deposition of collagen and presence of α-SMA-positive myofibroblasts. The authors also described elevated numbers of M2-type macrophages, which persisted in chronic lesions. Moreover, this study also analyzed the dynamics of AT2-derived ADI cells and demonstrated that persistence of ADI cells is a feature of chronic lesions, in line with our findings in the hamster model. However, the study did not investigate airway progenitor cell contribution to alveolar regeneration. In contrast to this, we found a prominent airway progenitor mobilization into damaged alveoli in our hamster model, indicating that hamsters model this aspect of lung regeneration observed in humans more closely than mice.

Although pre-existing AT2 cells are described to be the predominant source of AT1s after alveolar damage, it is known that other cell types partake in regenerative processes, especially after severe injury ^15,40^. In case of severe damage that involve broad epithelial denudation, basal cells can migrate into alveoli, become distal basal-like cells and subsequently promote alveolar regeneration giving rise to AT2 ^15,38,44^. A contribution of airway progenitors to alveolar repair has been reported in COVID-19 patients ^24,64^. In COVID-19 patients, the most prominent airway progenitors supporting alveolar regeneration were reported to be CK5^+^ basal cells, which form the so-called “keratin 5 pods”. Basal cell expansion, also termed “pod”“ is gradually recognized as common feature of epithelial remodeling ^17,44^. To a lesser extent, more immature CK5^+^p63^+^ basal cells were also reported to support alveolar regeneration in COVID-19 patients ^38^. Conversely, in SARS-CoV-2 infected hamsters, we found predominantly CK14^+^ cells within alveolar proliferation foci, resembling the human CK5^+^ pods. Basal airways cells originate from the same CK5^+^CK14^+^p63^+^ pool that gives rise to different combinations of CK5^+/-^, CK14^+/-^, p63^+/-^ progenitor cells that will populate the airways ^41^. Some of these cells also have the potential to give rise to AT2 cells ^40,41^. It appears that the subpopulation might differ among various species. Human lung multipotent cells can differ from murine ones, and in its turn, we might expect the same for other rodents like hamsters. The CK14^+^ basal cells detected in our study were having similar features like the ones described for human CK5^+^ cells, namely pods formation and differentiation towards AT2 cells, and therefore can be considered the hamster equivalent of human basal cells contributing to alveolar repair.

The authors recognize that the study has some limitations. First, this work provides a whole slide digital quantification of the main cell types involved in alveolar regeneration upon SARS-CoV-2 infection, including CK8^+^ ADI cells. However, since that several tested antibodies (anti-AGER, -AQP5, -PDPN) failed to specifically recognize AT1 cells in hamsters, a quantification of these cells and demonstration of ADI-AT1 transition by double-labeling was not possible. Therefore, the ADI-AT1 transition was demonstrated with ultrastructural analysis, in line with previous COVID-19 reports. Second, we can only speculate on the clinical relevance ADI persistence and fibrotic lesions in the animals. However, once that this work confirmed hamsters to be a reliable model for these features, further investigations including longer time points and the assessment of lung function and gas-exchange capacity are warranted. Third, the conclusions regarding cell origins in this work are based on double-labelling and co-expression of genes interpreted in the context of published literature. Additional studies involving lineage-tracing are required to irrefutably prove cell trajectories.

In conclusion, our study provides a detailed characterization of cell populations composing the pulmonary epithelial regenerative response in the hamster and thus provides preliminary and highly needed information about this important translational COVID-19 model. We show that ADI cells and airway-derived progenitors participate in alveolar regeneration in the species, and provide evidence of ongoing regeneration post virus-clearance. Thus, hamsters are a suitable model to investigate the relevance of these changes and their actual contribution to PASC symptoms. However, further studies including a longer investigation period and more detailed clinical analyses are required. Since post-COVID-19 pathological lesions show overlap with other diseases featuring DAD and IPF, the model can be used for broader implications.

## Material and methods

### Hamster study

The animal experiment was in accordance with the EU directive 2010/63/EU and approved by the relevant local authorities (protocol code N032/2020 22 April 2020). During the experiment the animals were under veterinary observation and all efforts were made to minimize distress. Eight to ten weeks old male and female Syrian golden hamsters (*Mesocricetus auratus*) purchased from Janvier Labs were housed under BSL-3 conditions for 2 weeks prior the experiment for acclimatization. A total of 80 hamsters divided into groups of 5 male and 5 female (n=10) animals per time point per infection group were housed in isolated ventilated cages under standardized conditions (21 ± 2 °C, 40 – 50 % relative humidity, 12:12 light-dark cycle, food and water ad libitum) at the Heinrich Pette Institute, Leibniz Institute for Experimental Virology in Hamburg, Germany. Animals were infected with an intranasal inoculation of either a suspension containing 10^5^ plaque-forming units (pfu) of SARS-CoV2 (SARS-CoV-2/Germany/Hamburg/01/2020; ENA study PRJEB41216 and sample ERS5312751) or phosphate-buffered saline (PBS, control) as previously described ^65^ under general anaesthesia. At 1, 3, 6 and 14 days post-infection (dpi), groups of five female and five male hamsters (n=10) per each treatment (either SARS-CoV-2 infected or mock infected) were euthanized by intraperitoneal administration of a pentobarbital-overdose and blood withdrawal by cardiac puncture. Immediately after death, right lung lobes (*lobus cranialis, lobus medius, lobus caudalis, lobus accessorius*) were collected and fixed in 10 % neutral-buffered formalin (Chemie Vetrieb GmbH & Co) or 5 % glutaraldehyde (Merck KGaA) for microscopic and ultrastructural evaluation respectively.

### Virus

SARS-CoV-2/Germany/Hamburg/01/2020 (ENA study PRJEB41216 and sample ERS5312751) was isolated from a nasopharyngeal swab of a confirmed COVID-19 patient. Stock virus was produced after three serial passages in Vero E6 cells using Dulbecco’s Modified Eagle’s Medium (DMEM; Sigma) supplemented with 2 % fetal bovine serum, 1 % penicillin-streptomycin and 1 % L-glutamine at 37 °C. The infection experiment was carried out under biosafety level 3 (BSL-3) conditions at the Heinrich Pette Institute, Leibniz Institute for Experimental Virology in Hamburg, Germany.

### Human samples

Lung samples were obtained from three patients who died of respiratory failure caused by severe COVID-19. The patients were two men, aged 76 and 74 years, and one woman, aged 74 years. The patients were hospitalized for 21, 7 and 5 days, respectively, and all received mechanical ventilation. SARS-CoV-2 infection was confirmed by PCR. The lung samples were obtained during autopsy. In addition, one non-COVID-19 lung sample was obtained from a 66-year-old man who underwent a lobectomy due to a pulmonary neoplasm. All patients or their relatives provided written informed consent for the use of their data and samples obtained during autopsy for scientific purposes. Ethical approval was given by the local institutional review board at Hannover Medical School (no. 9621_BO_K_2021).

### Histopathology

For histopathological evaluation, lung samples were formalin-fixed and embedded in paraffin. Serial sections of 2μm were cut and stained with hematoxylin and eosin (HE) and Azan trichrome. Qualitative evaluations with special emphasis on inflammatory and epithelial regenerative processes (HE) as well as on fibrosis (Azan) were performed in a blinded fashion by veterinary pathologists (FA, LH) and subsequently reviewed by board certified veterinary pathologist (MCI,WB).

### Immunohistochemistry

Immunohistochemistry was performed to detect SARS-CoV-2 antigen (SARS-CoV-2 nucleo protein), macrophages and dendritic cells (ionized calcium-binding adapter molecule 1, IBA-1), alveolar pneumocytes type 2 (pro surfactant protein C), alveolar differentiation intermediate cells (cytokeratin 8), airway basal cells (cytokeratin 14), club cells (secretoglobin 1A1), and M2 macrophages (CD 204). Immunolabelings were visualized either using the Dako EnVision+ polymer system (Dako Agilent Pathology Solutions) and 3,3’-Diaminobenzidine tetrahydrochloride (DAB, Carl Roth) as previously described ^34^ or using avidin–biotin complex (ABC) peroxidase kit (Vector Labs) and DAB (Carl Roth) as previously described ^66^. Nuclei were counterstained with hematoxylin. Further details about primary and secondary antibodies, visualization methods and dilutions used can be found in supplementary table 2. For negative controls, the primary antibodies were replaced with rabbit serum or BALB/cJ mouse ascitic fluid, respectively, with the dilution chosen according to protein concentration of the exchanged primary antibody. Antibodies were tested on murine and human lung tissue to confirm specificity for the cells of interest. Subsequently, murine and human tissues were used as positive controls.

### Immunofluorescence

Double labelling immunofluorescence was performed to investigate different states of alveolar pneumocytes type 2 and alveolar diffentiation intermediate cells, as well as to prove that airways progenitor cells can differentiate into alveolar cell types. Reaction was carried out as previously described with minor modifications ^67^. Briefly, after deparaffinization, HIER and serum blocking, washing with PBS in between each step, a dilution containing two primary antibodies was added and incubated overnight at 4 °C. Afterwards, a dilution containing two secondary antibodies were incubated for 60 minutes at room temperature in the dark. After washing with PBS and distilled water, sections were counterstained and mounted using anti-fade mounting medium containing DAPI (Vectashield®HardSet™, Biozol). Further details about primary and secondary antibodies, visualization methods and dilutions used can be found in in supplementary table 3. For negative controls, the primary antibodies were replaced with rabbit serum or BALB/cJ mouse ascitic fluid respectively with the dilution chosen according to protein concentration of the exchanged primary antibody.

### Transmission Electron Microscopy (TEM)

In order to detect AT1 cells with features of AT2 proving the final trajectory ADI-AT1 in hamsters, transmission electron microscopy was performed. Reactions were carried out as previously described ^65,68^. Briefly, glutaraldehyde-fixed lung tissue was rinsed overnight in cacodylate buffer (Serva Electrophoresis GmbH), followed by post-fixation treatment in 1 % osmium tetroxode (Roth C. GmbH & Co. KG). After dehydration using a graded alcohol series, samples were embedded in epoxy resin. Representative areas of affected alveoli were then cut into ultrathin sections, contrasted with uranyl acetate and lead acetate and subsequently morphologically evaluated using a transmission electron microscope (EM 10C, Carl Zeiss Microscopy GmbH).

### Digital image analysis

To quantify immunolabeled cells in pulmonary tissue, areas of alveolar epithelial proliferation as well as areas of subpleural fibrosis, slides were digitized using an Olympus VS200 Digital slide scanner (Olympus Deutschland GmbH). Image analysis was performed using QuPath (version 0.3.1), an open-source software package for digital pathology image analysis ^69^. For all animals, whole slide images of the entire right lung were evaluated. or the pro surfactant protein C (proSPC), cytokeratin 8 (CK8), cytokeratin 14 (CK14), secretoglobin 1A1 (SCGB1A1) immunolabelings, total lung tissue was first detected automatically using digital thresholding. Afterwards, regions of interest (ROI) were defined. The ROIs “airways” (bronchi, bronchioli, terminal bronchioli), “blood vessels”, “affected alveoli” (alveoli that were involved either in an inflammatory process or in a epithelial regenerative process or both) and “artifacts” were manually outlined. The area denoted as “total alveoli” was defined by subtraction of the “blood vessels”, “airways” and “artifacts” ROIs from the total lung tissue using an automated script. The area denoted as “unaffected alveoli” (alveoli that were morphologically free from any inflammatory or regenerative process) was defined by subtracting the ROI “affected alveoli” from the ROI “total alveoli” using an automated script. Using tissue- and marker-specific thresholding parameters, quantification of immunolabeled cells was achieved by automated positive cell detection in all ROI. To analyze SARS-CoV-2 NP, IBA-1and CD204 immunolabeling, total lung tissue was automatically detected using digital thresholding. Afterwards, only blood vessels and artifacts were indicated as ROIs and subtracted from the total lung tissue. Based on tissue and marker specific thresholding parameters, quantification of immunolabeled cells was then achieved by automated positive cell detection. For quantification of alveolar epithelial proliferation or subpleural fibrosis, total lung tissue area was automatically detected using digital thresholding. Subsequently, either alveolar epithelial proliferation or subpleural fibrosis were marked as ROIs and the total area was calculated. Finally, the percentage of total lung area affected by either epithelial proliferation or subpleural fibrosis was obtained. All procedures (tissue detection, indication of ROIs, positive cell detection) were performed and subsequently reviewed by at least two veterinary pathologists (FA, GB, LH, MC). Statistical analysis and graphs design were performed using GraphPad Prism 9.3.1 (GraphPad Software, San Diego, CA, USA) for Windows™. Single comparison between SARS-CoV-2 infected hamsters and control group were tested with a two-tailed Mann–Whitney-U test. For multiple comparisons among different time-points data were tested for significant differences using Kruskal–Wallis tests and corrected for multiple group comparisons using the Benjamini–Hochberg correction. Statistical significance was accepted at exact p-values of ≤0.05.

### single-cell RNAseq

Single-cell RNASeq data from lungs of SARS-CoV-2 infected hamsters was obtained from a publicly available dataset ^43^. Data were analyzed using the R software package (version 3.6.0) ^70^. Expression data were downloaded from GEO (https://www.ncbi.nlm.nih.gov/geo/, GSE162208) and Seurat objects (version Seurat_3.2.0, ^71-74^ were generated from h5 files by combining replicate samples from lung day5 and day14. Pre-processing of data was performed by applying several Seurat functions: subset (subset = nFeature_RNA > 200 & nFeature_RNA < 2500 & percent.mt < 5), NormalizeData (data), FindVariableFeatures (data, selection.method = “vst”, nfeatures = 2000), ScaleData(data, features = all.genes), and clusters identified using functions RunPCA(data, features = VariableFeatures (object = data)), FindNeighbors (data, dims = 1:10), FindClusters (data, resolution = 0.5). AT1 and AT2 cell cluster were then identified by using the marker genes *Rtkn2* (AT1) and *Lamp3* (AT2), respectively, from the original publication ^43^. These clusters were selected, then pre-processed and re-clustered as described above. We then collected more candidate marker genes for AT1, AT2 and additional cell populations in these clusters by applying functions FindAllMarkers (pbmc, only.pos = TRUE, min.pct = 0.25, logfc.threshold = 0.25) and selecting the top 10 markers genes per cluster. We further identified additional candidate markers from ^9,19,43^. We evaluated the specificity of these candidate markers by visualizing them with the functions FeaturePlot, DoHeatmap and AddModuleScore. The list of final maintained marker genes is presented in supplementary table 1. The function AddModuleScore was then used to visualize the various cell populations and hallmark genes from the GSEA database (^75^ http://www.gsea-msigdb.org/gsea/msigdb).

## Data Availability

Source data will be provided with this paper.

## Acknowledgements

The authors are grateful to Julia Baskas, Petra Grünig, Jana-Svea Harre, Kerstin Rohn, Caroline Schütz, and Kerstin Schöne for excellent technical assistance. This project was in part supported by the COVID-19 Research Network of the State of Lower Saxony (COFONI) with funding from the Ministry of Science and Culture of Lower Saxony, Germany (14-76403-184, project number 5FF22, Federico Armando, Wolfgang Baumgärtner, Malgorzata Ciurkiewicz). This research was in part supported by the Deutsche Forschungsgemeinschaft (DFG; German Research Foundation) -398066876/GRK 2485/1, Wolfgang Baumgärtner, Georg Beythien, Laura Heydemann). This study was also supported in part by intra-mural grants from the Helmholtz-Association (Program Infection and Immunity), and NIAID Research Grants 2-U19-AI100625-06 REVISED and 5U19A|100625-07 awarded to Klaus Schughart.

## Author Contributions Statement

The study was designed by FA, WB and MC. The animal experiments were performed by SS-B, BS, NM-K, SB, MZ and GG. Histology, immunolabelling and electron microscopy evaluation of hamster tissues was conducted and analyzed by FA, LH, MC, GB, KB, AB and WB. Pathological analysis of human samples was performed by MK. scRNA-seq analysis was performed by KS. Data analysis and interpretation were performed by FA, LH, MC and GB. Figures were prepared by MC, KS and FA. The original draft was written by LH, MC, FA and KS. The manuscript was reviewed, edited, and approved by all authors. Funding was acquired by MC, KS and WB. The project was supervised by WB and FA

## SUPPLEMENTARY DATA

**Supplementary table 1:** Marker gene list of main pulmonary cell populations in the hamster species.

| AT2 | AT1 | ADI | CLUB | CILIATED |
| --- | --- | --- | --- | --- |
| *Sftpc* | *Col12a1* | *S100a10* | *Pigr* | *Ccdc153* |
| *Napsa* | *Sema3a* | *Anxa3* | *Ifitm2* | *Caps* |
| *Fabp5* | *Dag1* | *S100a6* | *Gss* | *Dynlrb2* |
| *Scd1* | *Aqp5* | *Krt7* | *Scgb3a2* | *1700094D03Rik* |
| *Sftpa1* | *Gprc5a* | *Krt8* | *Hp* |  |
| *Ctsc* | *Cav1* | *Dstn* | *Fam216b* |  |
| *Lamp3* | *Vegfa* | *Anxa1* |  |  |
| *Pgc* | *Cav2* | *Tuft1* |  |  |
| *Lgi3* | *Itga3* | *Tacstd2* |  |  |
| *Sfta2* | *Lama5* | *Klf6* |  |  |
| *Gas6* | *Nckap5* | *Lmo7* |  |  |
| *Egfl6* | *Abca5* | *Cdkn1a* |  |  |
| *Lcn2* | *Itm2a* | *Tp53* |  |  |
| *Atp1a1* | *Limd2* | *Tnip3* |  |  |
| *Abca3* | *Wsb1* | *Hbegf* |  |  |
| *Sftpd* | *Sec14l3* | *Ggh* |  |  |
| *Slc34a2* | *Prdx6* | *Steap4* |  |  |
| *Serpine2* | *Mfge8* | *Zfp36* |  |  |
|  | *Ccnd2* | *Junb* |  |  |
|  | *Timp3* | *Jun* |  |  |
|  |  | *Fos* |  |  |
|  |  | *CRYAB* |  |  |
|  |  | *Ndnf* |  |  |
|  |  | *Timp2* |  |  |
|  |  | *Emp2* |  |  |
|  |  | *Sox4* |  |  |
|  |  | *Wwtr1* |  |  |
|  |  | *Sparc* |  |  |

Abbreviations: alveolar pneumocytes type 2 (AT2), alveolar pneumocytes type 1 (AT1), alveolar differentiation intermediate cells (ADI), club cells (CLUB), ciliated cells (CILIATED)

**Supplementary table 2:** Primary antibodies, visualization method, dilution, clonality and host species, secondary antibody as well as positive controls used for immunohistochemical investigations.

| Primary antibody | Visualization method | Dilution | Clonality, host species | Secondary antibody (1:200) | Positive control |
| --- | --- | --- | --- | --- | --- |
| CK8 (Invitrogen, PA5-29607) | EnVision | 1 : 250 | Polyclonal, rabbit | / | Airways (Hu, Ms, Hm internal control) |
| CK14 (Invitrogen, PA5-16722) | ABC | 1 : 500 | Polyclonal, rabbit | GAR-b | Airways (Hu, Ms, Hm internal control) |
| SCGB1A1 (Proteintec, 10490-1-AP) | ABC | 1 : 200 | Polyclonal, rabbit | GAR-b | Airways (Hu, Ms, Hm internal control) |
| proSP-C (MEMD Millipore, AB3786 ) | EnVision | 1 : 1000 | Polyclonal, rabbit | / | Alveolar AT2 (Hu, Ms, Hm internal control) |
| IBA-1 (FUJIFILM Wako Pure Chemical Corporation, 019–19741) | ABC | 1 : 500 | Polyclonal, rabbit | GAR-b | SARS-CoV-2 Infected lung, 6 dpi (Hm internal control) |
| SARS CoV-2-NP (Sinobiological, 40143-MM05) | EnVision | 1 : 16000 | Monoclonal, mouse, clone 5 | GAM-b | SARS-CoV-2 Infected lung, 3 dpi (Hm internal control) |
| α-SMA (Dako, GA611) | ABC | 1: 500 | Monoclonal, mouse, Clone 1A4 | GAM-b | Airways and vessels (Hu, Ms, Hm internal control) |
| CD-204 (Abnova Corporation, MAB1710) | ABC | 1:1000 | Monoclonal, mouse, clone SRA-E5 | GAM-b | SARS-CoV-2 Infected lung, 6 dpi (Hm internal control) |

Abbreviations: α-SMA: alpha smooth muscle actin; ABC: Avidin-biotin-complex; CK8: cytokeratin 8; CK14: cytokeratin 14; GAR-b: goat anti rabbit-biotinylated; GAM-b: goat anti mouse-biotinylated; Hm: hamster; Hu: human; IBA-1: ionized calcium-binding adapter molecule 1; Ms: mouse; SARS-CoV-2 NP: severe acute respiratory syndrome coronavirus-2 nucleoprotein; SCGB1A1: secretoglobin 1A1; proSP-c: pro surfactant protein C;

**Supplementary table 3:** Primary antibodies, dilution, clonality and host species as well as secondary antibody used for immunofluorescence investigations.

| Primary antibody | Dilution | Clonality, host species | Secondary antibody<br>(1:200) |
| --- | --- | --- | --- |
| CK8 (Invitrogen, PA5-29607) | 1 : 500 | Polyclonal, rabbit | GAR Cy2/ GAR Cy3 |
| CK8-FITC conjugated (abcam, ab192467) | 1 : 200 | Monoclonal, rabbit, clone EP1628Y | / |
| CK14 (Invitrogen, PA5-16722) | 1 : 500 | Polyclonal, rabbit | GAR Cy2 |
| CK14 (Invitrogen, MA5-11599) | 1 : 500 | Monoclonal, mouse, clone LL002 | GAM Cy2/ GAMCy3 |
| proSP-C (MEMD Millipore, AB3786 ) | 1 : 1000 | Polyclonal, rabbit | GAR Cy3 |
| SCGB1A1 (Proteintec, 10490-1-AP) | 1 : 200 | Polyclonal, rabbit | GAR Cy3 |
| TP53 (Novusbio, NBP2-29453) | 1 : 100 | Monoclonal, mouse, clone BP53-12 | GAM Cy3 |
| ΔNp63 (Cell signalling, #67825S) | 1 : 800 | Monoclonal, rabbit, clone E6Q30 | GAR Cy3 |
| CK5-FITC conjugated (Abcam, ab-193894) | 1 : 200 | Monoclonal, rabbit, clone EP1601Y | / |

Abbreviations: Cy2: cyanin 2 conjugated; Cy3: cyanin 3 conjugated; CK8: cytokeratin 8; CK14: cytokeratin 14; CK5: cytokeratin 5; GAR-b: goat anti rabbit-biotinylated; GAM-b: goat anti mouse-biotinylated;

**Supplementary figure 1:**
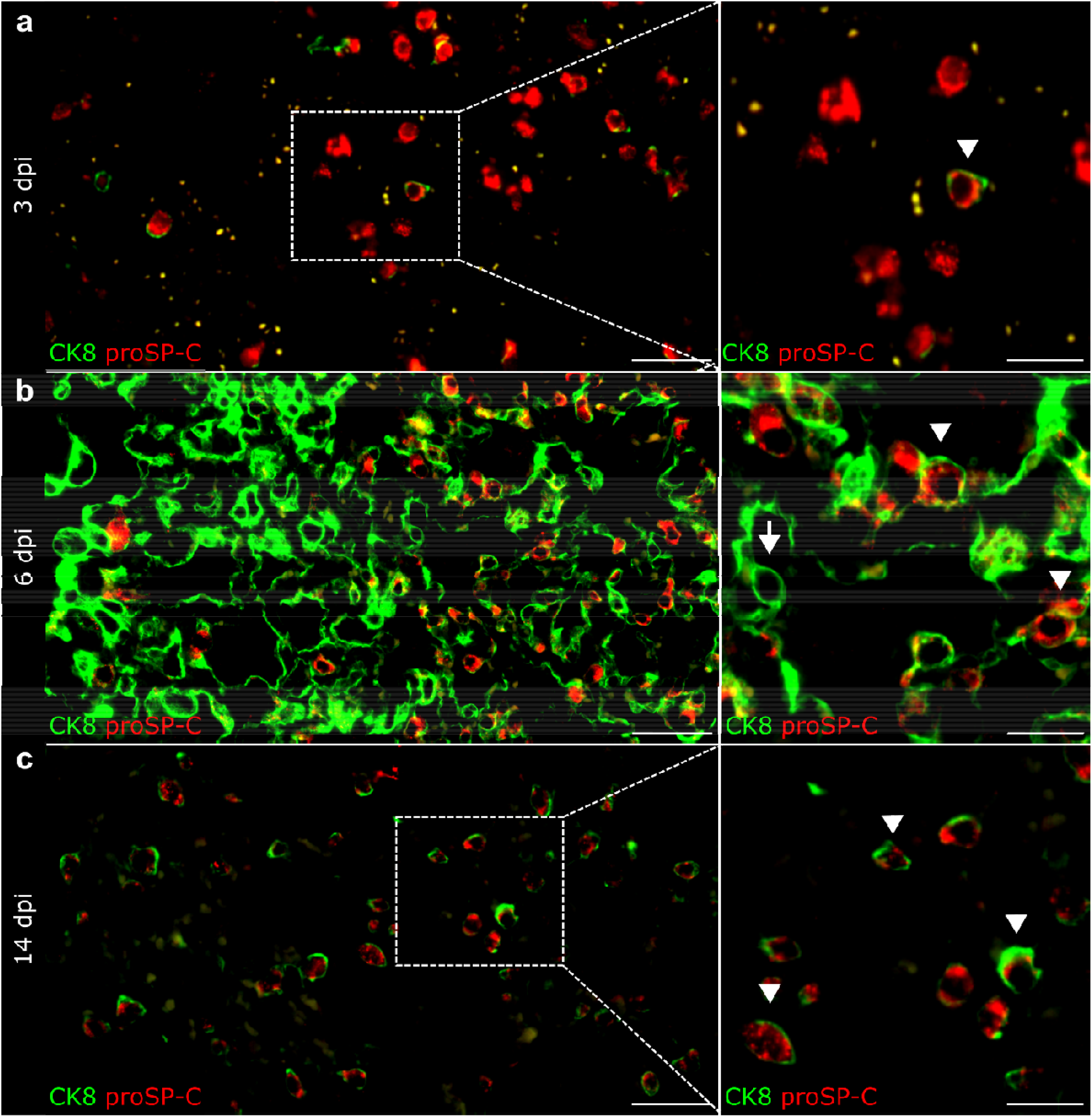
Alveolar pneumocytes type 2 (AT2) - alveolar differentiation intermediate (ADI) cell trajectory in SARS-CoV-2 infected hamsters at different time points post infection. Representative double immunofluorescence images of alveoli in a SARS-CoV-2 infected hamsters at 3 (A), 6 (B) and 14 (C) days post infection (dpi). Cells are labelled with CK8 (green) and proSP-C (red). For each time point, an overview and higher magnification of the area delineated by the rectangle are shown. **A** At 3 dpi, there are numerous, round, proSP-C^+^CK8^-^ AT2 cells and rare, round, proSP-C^+^CK8^+^ ADI cells (arrowhead). **B** Alveolar proliferation at 6 dpi contain numerous proSP-C^-^ CK8^+^ADI cells, some showing hypertrophy and elongated cytoplasmic processes (arrow). There are single proSP-C^+^CK8^+^ cells with a round morphology (arrowheads). **C** At 14 dpi, there are numerous round, proSP-C^+^CK8^+^ ADI cells (arrowheads) and rare round proSP-C^+^CK8^-^ AT2 cells. Scale bars: 50 µm (left panel) and 20 µm (right panel).

**Supplementary figure 2:**
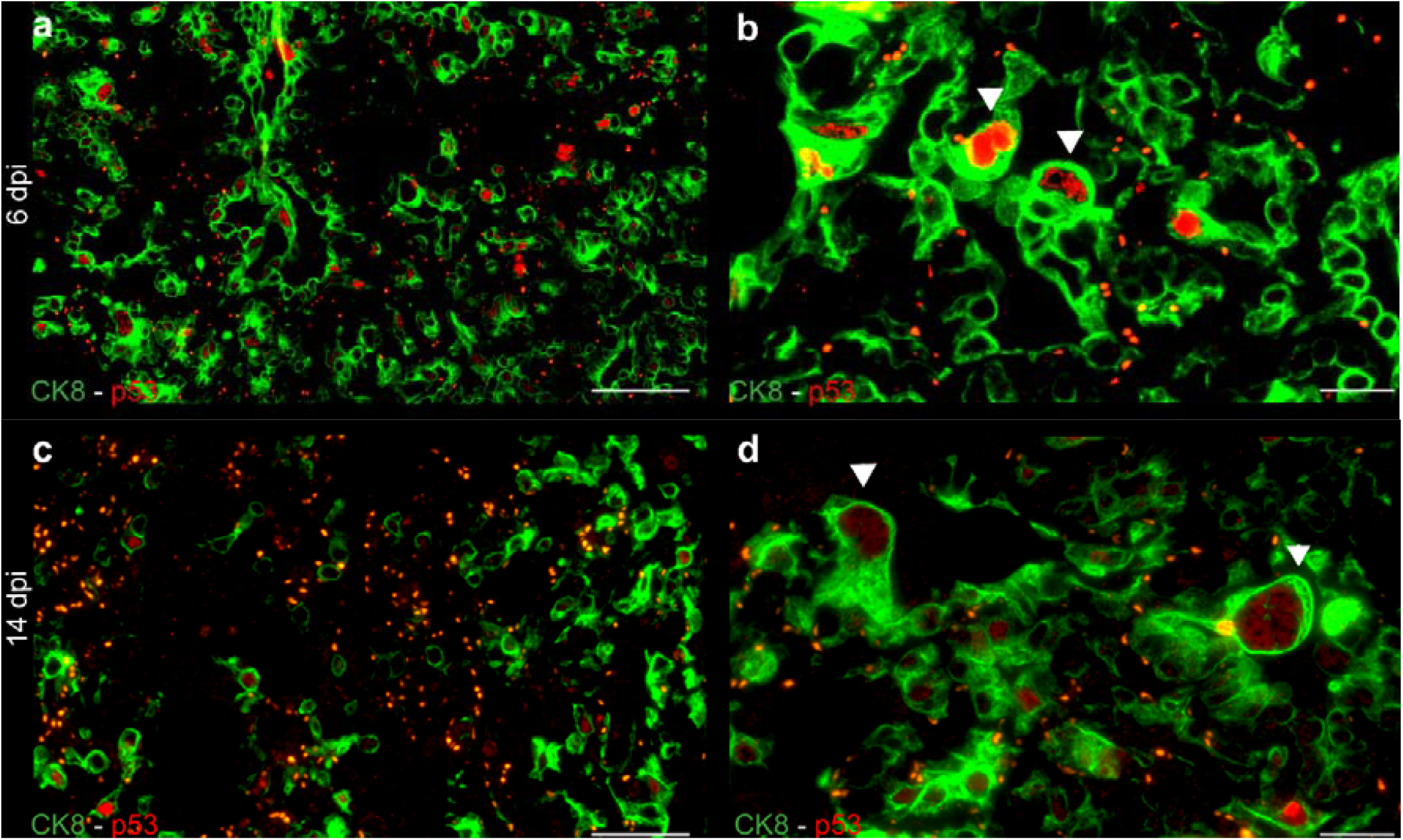
Alveolar differentiation intermediate (ADI) cells exhibit cell cycle arrest in SARS-CoV-2 infected hamsters at different time points post infection. Representative double immunofluorescence images of alveoli in SARS-CoV-2 infected hamsters at 6 (A, B) and 14 (C, D) days post infection (dpi). Cells are labelled with CK8 (green) and TP53 (red). **A, B** Overview and high magnification of proliferation focus at 6 dpi showing numerous CK8^+^ ADI expressing nuclear TP53. The high magnification shows polygonal, large, bizarre TP53^+^ ADI cells (arrowheads). **C** Overview of morphologically normal alveoli at 14 dpi showing numerous TP53^+^ ADI cells with a round morphology. **D** Large, bizarre TP53^+^ ADI cells (arrowheads) within residual alveolar lesions at 14 dpi. Scale bars: 50 µm and 20 µm (b,d).

**Supplementary figure 3:**
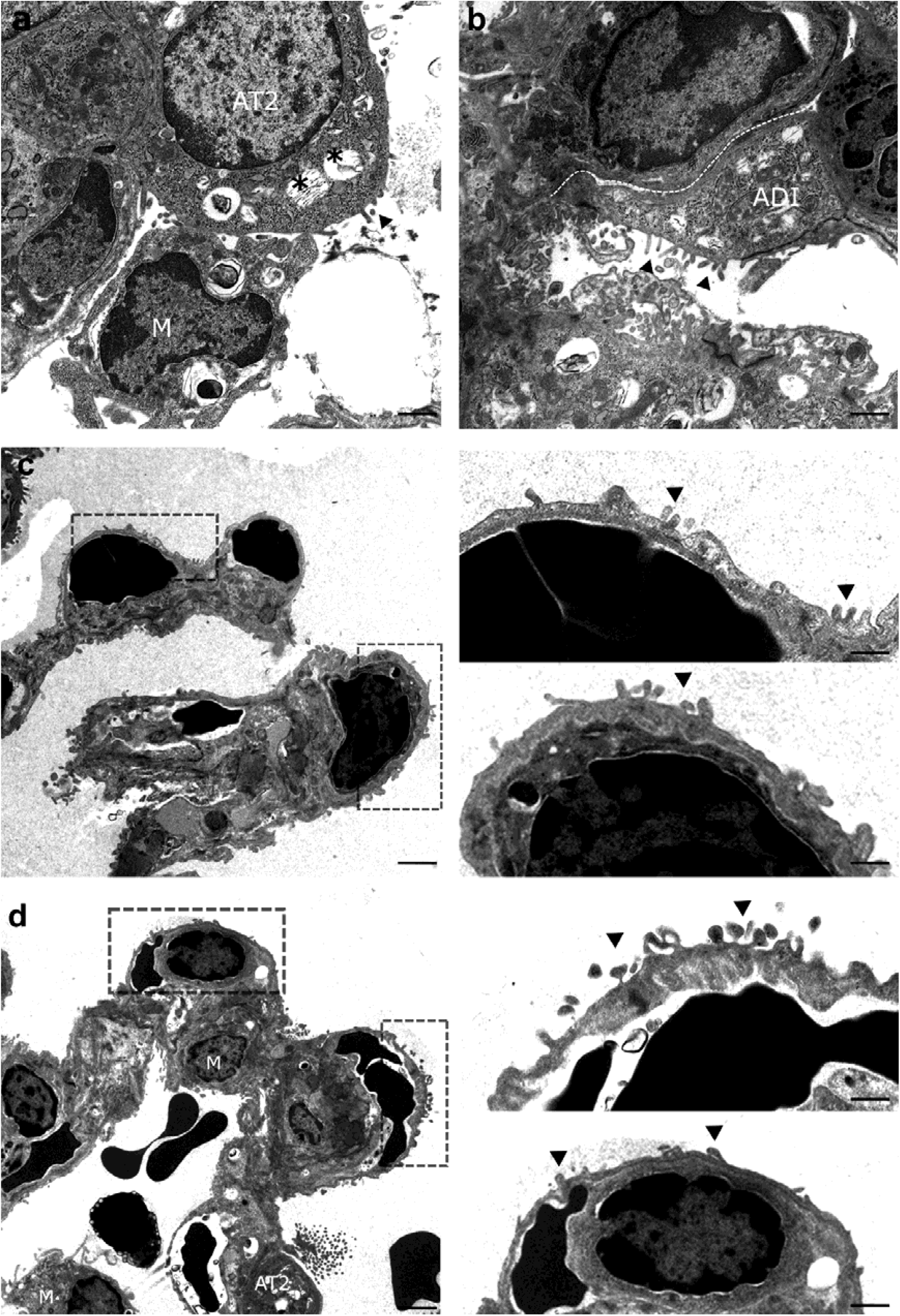
Alveolar pneumocytes type 1 (AT1) - alveolar differentiation intermediate (ADI) cells trajectory in SARS-CoV-2 infected hamsters. A Transmission electron microscopy (TEM) micrograph of normal alveolar cells showing a round cell (AT2) with apico-basal polarity, apical microvilli (arrowhead), moderately electron-dense cytoplasm, rich in rough endoplasmic reticulum and free ribosomes as well as numerous membrane-bound vesicles containing multiple concentric membrane layers (multi-lamellar bodies, asterisks). On the left, a macrophage with intracytoplasmic multi-lamellar bodies is also seen (M). **B** Representative micrograph showing alveoli of a SARS-CoV-2 infected hamster at 6 dpi. In the center, a stretching cell (ADI) showing AT2 features like ribosome-rich cytoplasm and microvilli (arrowheads) on the cell surface is seen. The dotted line indicates the basal contour of the cell, highlighting the elongated shape. **C, D** Representative micrograph showing alveoli of SARS-CoV-2 infected hamsters at 6 dpi. Overviews and higher magnification of the areas delineated by rectangles are shown. Alveolar septae are covered by delicate, elongated cells, separated by a thin basement membrane from capillaries containing erythrocytes. Macrophages (M) as well as an AT2 cell (AT2) are also seen. Rectangles and high magnification show cells with flattened and elongated morphology of AT1 cells but also retained characteristics of AT2 cells, such as apical microvilli (arrowheads), indicative of a transitional state. Scale bars: 1000 nm (a), 500 nm, 2000 (overview in c, d), 500 nm (high magnifications in c, d).

**Supplementary figure 4:**
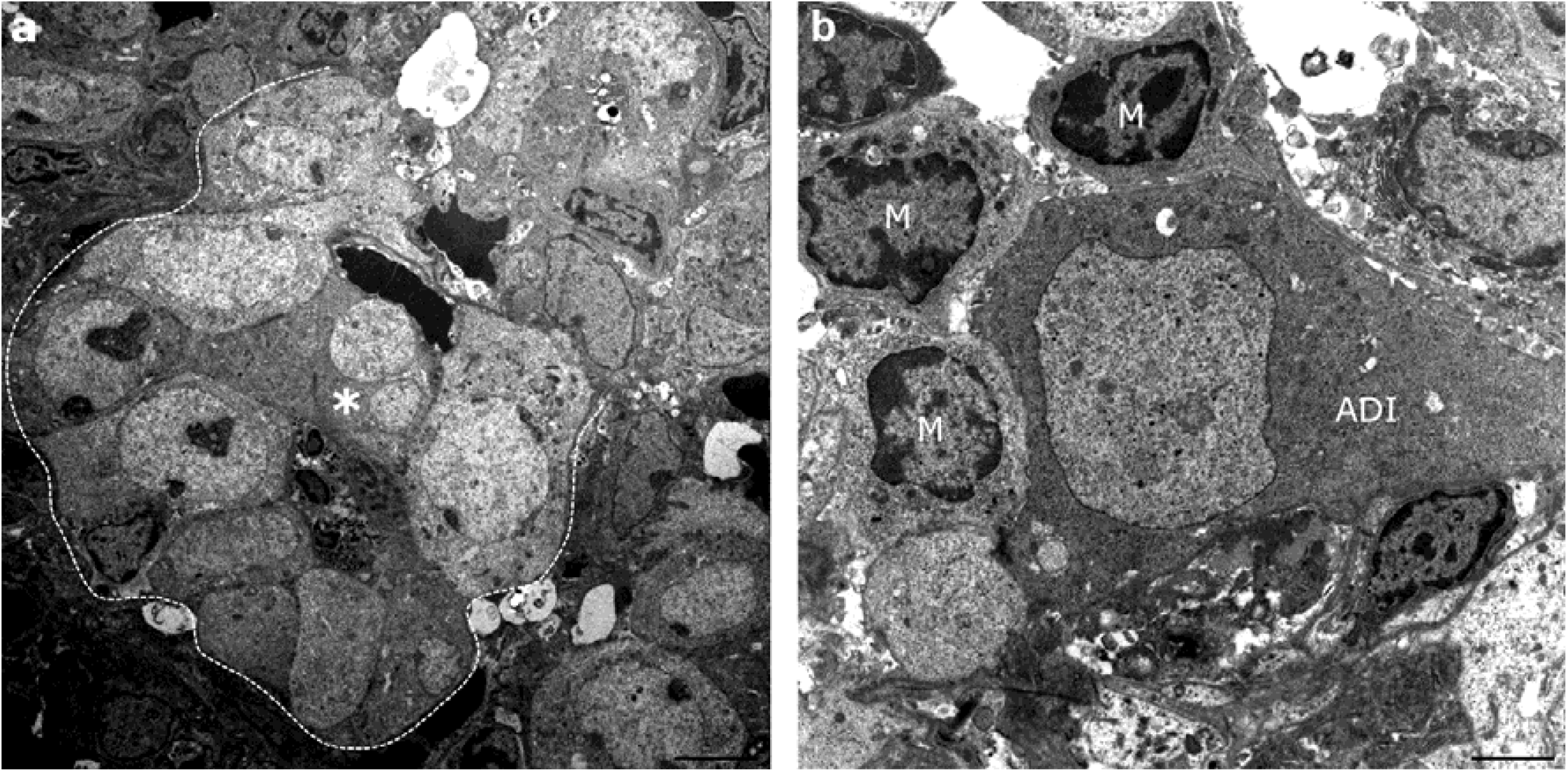
Epithelial proliferates and in SARS-CoV-2 infected hamsters. A Transmission electron microscopy (TEM) micrograph of an epithelial proliferation focus from a SARS-CoV-2 infected hamster at 6 days post infection (dpi). A string of proliferating, polygonal to columnar epithelial cells forming a tubule-like structure (dotted line) is shown. The cells show hypertrophy, numerous prominent mitochondria and irregularly clumped chromatin. A bizarre cell with two unevenly large nuclei is also present (asterisk). **B** Micrograph of an ADI cell within a proliferation focus from a SARS-CoV-2 infected hamster at 6 dpi. A triangular, hypertrophic ADI cell surrounded by multiple macrophages (M) is shown. Scale bars: 500 nm.

**Supplementary figure 5:**
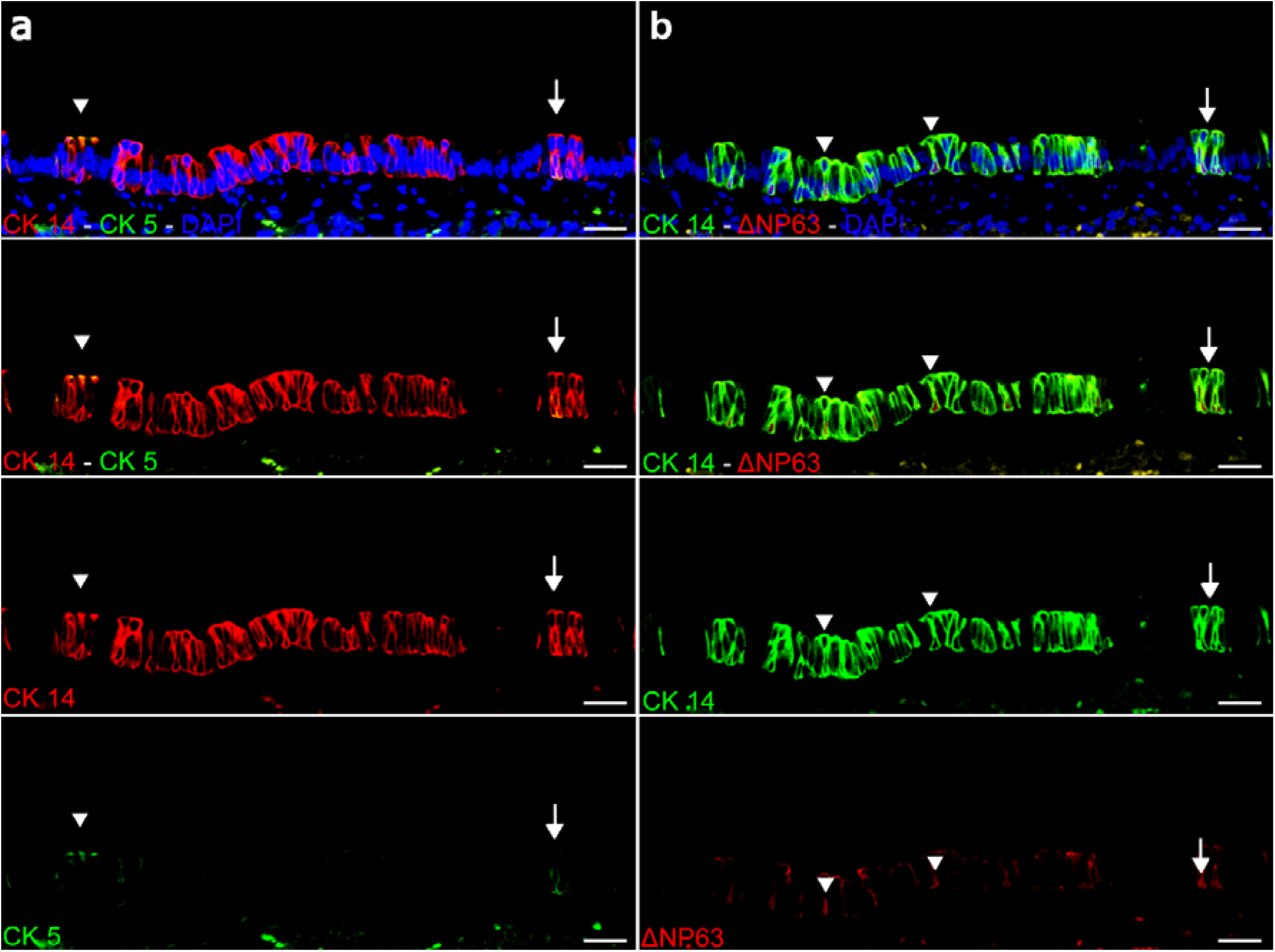
basal cells in the airways of hamsters. A Representative images of double immunofluorescence for CK5 (green) and CK14 (red) in a hamster airway, showing numerous CK14^+^CK5^-^ cells and occasional double labelled CK14^+^CK5^+^ cells (arrowheads). **B** Representative image of double immunofluorescence for CK14 (green) and ΔNP63 (red) in a hamster airway, taken at the same location as the images in a. There are numerous CK14^+^ΔNp63^-^ cells and fewer CK14^+^ΔNp63^+^ cells (arrowheads). Cells indicated by an arrow in A and B are considered CK5^+^CK14^+^ΔNp63^+^. Scale bars: 25 µm.

**Supplementary figure 6:**
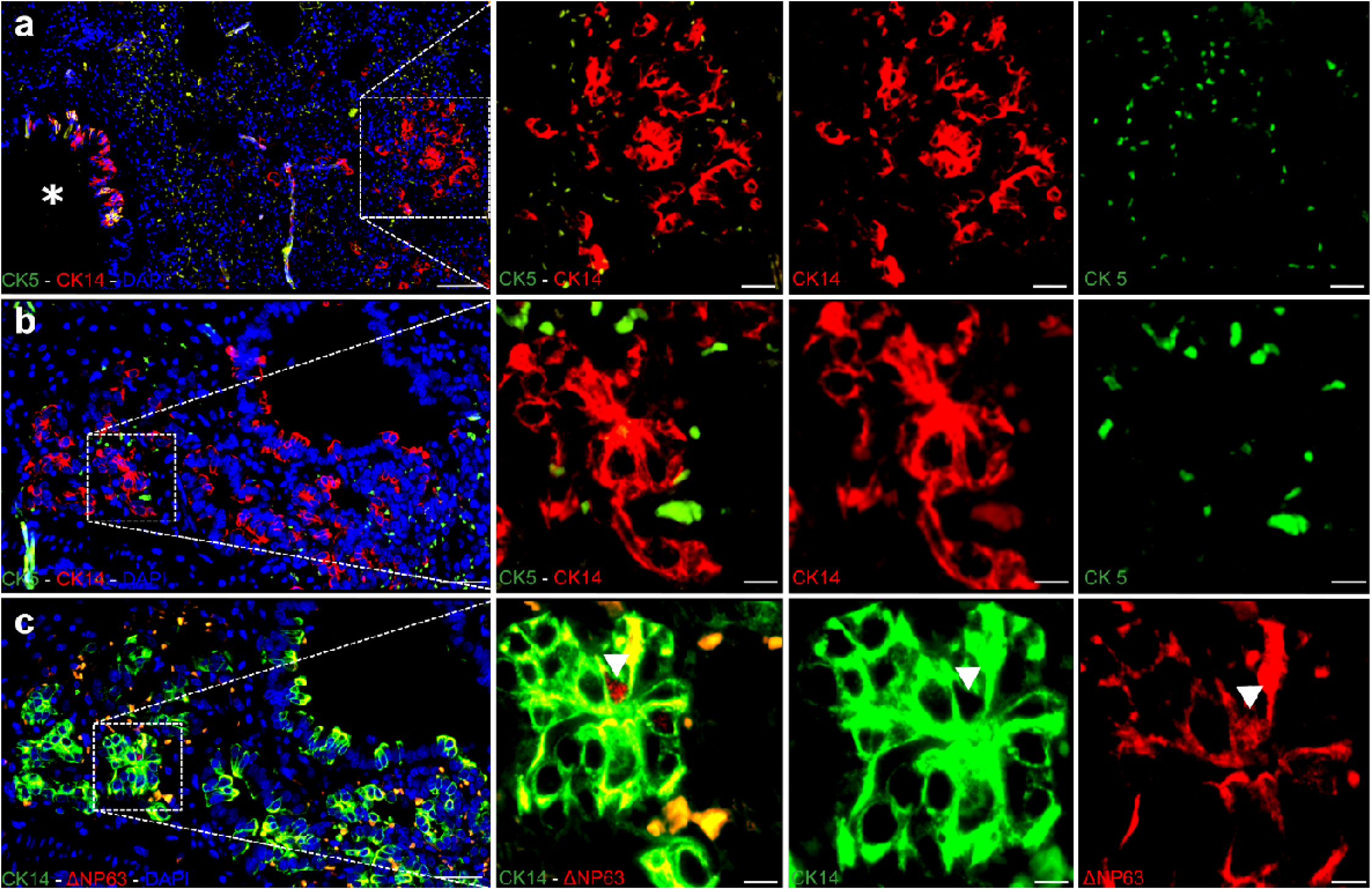
CK14^+^ and CK14^+^ΔNp63^+^ basal cells take part in alveolar proliferation in SARS-CoV-2 infected hamsters. A Representative image of double immunofluorescence for CK5 (green) and CK14 (red) in a peribronchiolar proliferation area in a SARS-CoV-2 infected hamster at 6 dpi. An overview and higher magnification of the area delineated by a rectangle are shown. The asterisk indicates a bronchiole containing CK5^+^CK14^+^ basal cells. Alveolar proliferation foci are composed of CK5^-^CK14^+^ basal cells. **B, C** Representative images of double immunofluorescence for CK5 (green) and CK14 (red) as well as CK14 (green) and ΔNP63 (red), respectively. Pictures are taken from the same peribronchiolar proliferation area in a SARS-CoV-2 infected hamster at 6 dpi. Overviews and higher magnification of the area delineated by rectangles are shown. Alveolar proliferation foci are mainly composed of CK5^-^CK14^+^ΔNP63^-^ and rare CK5^-^CK14^+^ΔNP63^+^ basal cells (arrowhead). Scale bars: 50 µm (overviews), 25 µm (high magnifications in a), 10 µm (high magnifications in b, c).

**Supplementary figure 7:**
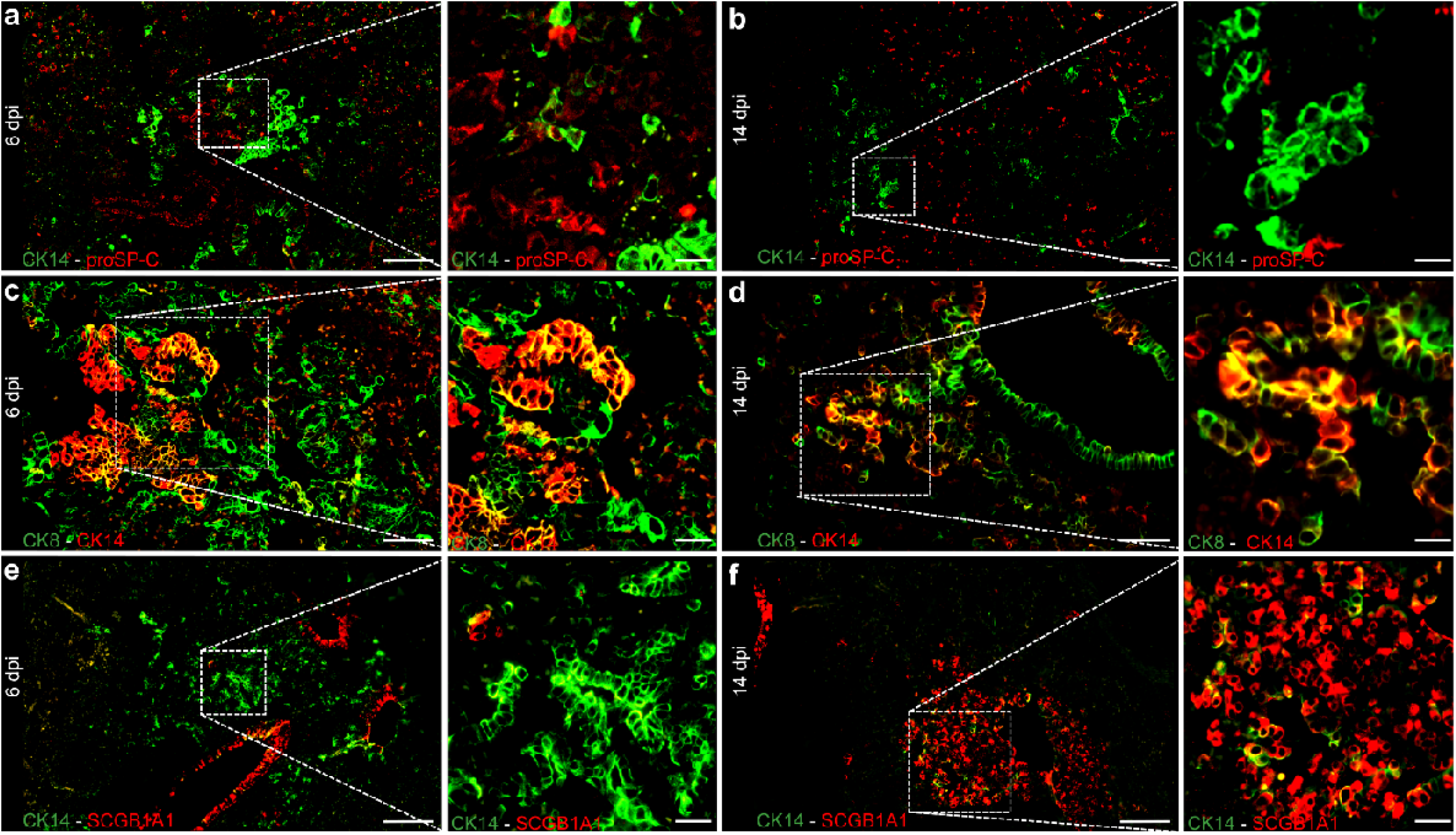
Airway basal cells in alveolar proliferates of SARS-CoV-2 infected hamsters at different time points. **A, B** Representative images of double immunofluorescence for CK14 (green) and proSP-C (red) in peribronchiolar proliferation areas in SARS-CoV-2 infected hamsters at 6 and 14 dpi, respectively. An overview and higher magnification of the areas delineated by rectangles are shown. **A** CK14^+^ basal cells in the alveoli. Single cells are CK14^+^proSP-C^+^, indicating that airway progenitors differentiate into proSP-C^+^ AT2 cells at 6 dpi. **B** CK14^+^ basal cells populating the alveolar proliferation foci without transdifferentiating in AT2 cells at 14 dpi. **C, D** Representative images of double immunofluorescence for CK14 (red) and CK8 (green) in peribronchiolar proliferation areas in SARS-CoV-2 infected hamsters at 6 and 14 dpi, respectively. An overview and higher magnification of the area delineated by a rectangle are shown. **C** Transition from CK14^+^ airway basal cells forming a pod, to double labeled CK14^+^CK8^+^ cells differentiating into CK14^-^CK8^+^, elongated ADI cells at 6 dpi. **D** CK14^+^ cells with airway-like morphology, populating the alveolar proliferation foci without transdifferentiating in elongated ADI cells at 14 dpi. **E, F** Representative images of double immunofluorescence for CK14 (green) and SCGB1A1 (red) in peribronchiolar proliferation areas in SARS-CoV-2 infected hamsters at 6 and 14 dpi, respectively. An overview and higher magnification of the area delineated by a rectangle are shown. **E** CK14^+^ cells populating the alveolar proliferatio focis without transdifferentiating in SCGB1A1^+^ club cells at 6 dpi. **F** CK14^+^ basal cells in the alveoli. Numerous cells are CK14^+^SCGB1A1^+^ indicating that airway progenitors differentiate into SCGB1A1^+^ club cells at 14 dpi. Scale bars: 50 µm (overviews), 25 µm (high magnifications).

## REFERENCES

1 WHO. A clinical case definition of post COVID-19 condition by a Delphi consensus, 6 October 2021, <https://www.who.int/publications/i/item/WHO-2019-nCoV-Post_COVID-19_condition-Clinical_case_definition-2021.1> (2021).

2 Nalbandian, A. et al. Post-acute COVID-19 syndrome. Nat. Med. 27, 601–615, doi:10.1038/s41591-021-01283-z (2021).

3 Castanares-Zapatero, D. et al. Pathophysiology and mechanism of long COVID: a comprehensive review. Ann Med 54, 1473–1487, doi:10.1080/07853890.2022.2076901 (2022).

4 Sudre, C. H. et al. Attributes and predictors of long COVID. Nat Med 27, 626–631, doi:10.1038/s41591-021-01292-y (2021).

5 Ayoubkhani, D. P. & Gaughan, C. Technical article: Updated estimates of the prevalence of post-acute symptoms among people with coronavirus (COVID-19) in the UK: 26 April 2020 to 1 August 2021., <https://www.ons.gov.uk/peoplepopulationandcommunity/healthandsocialcare/conditionsanddiseases/articles/technicalarticleupdatedestimatesoftheprevalenceofpostacutesymptomsamongpeoplewithcoronaviruscovid19intheuk/26april2020to1august2021> (2021).

6 Staudt, A. et al. Associations of Post-Acute COVID syndrome with physiological and clinical measures 10 months after hospitalization in patients of the first wave. Eur J Intern Med 95, 50–60, doi:10.1016/j.ejim.2021.10.031 (2022).

7 Fernandez-de-Las-Penas, C., Martin-Guerrero, J. D., Cancela-Cilleruelo, I., Moro-Lopez-Menchero, P. & Pellicer-Valero, O. J. Exploring the recovery curve for long-term post-COVID dyspnea and fatigue. Eur J Intern Med 101, 120–123, doi:10.1016/j.ejim.2022.03.036 (2022).

8 Ting, C. et al. Fatal COVID-19 and Non-COVID-19 Acute Respiratory Distress Syndrome Is Associated with Incomplete Alveolar Type 1 Epithelial Cell Differentiation from the Transitional State without Fibrosis. Am J Pathol 192, 454–467, doi:10.1016/j.ajpath.2021.11.014 (2022).

9 Melms, J. C. et al. A molecular single-cell lung atlas of lethal COVID-19. Nature 595, 114–119, doi:10.1038/s41586-021-03569-1 (2021).

10 Parimon, T. et al. Potential Mechanisms for Lung Fibrosis Associated with COVID-19 Infection. QJM, doi:10.1093/qjmed/hcac206 (2022).

11 Carsana, L. et al. Pulmonary post-mortem findings in a series of COVID-19 cases from northern Italy: a two-centre descriptive study. The Lancet Infectious Diseases 20, 1135–1140, doi:10.1016/s1473-3099(20)30434-5 (2020).

12 Polak, S. B., Van Gool, I. C., Cohen, D., von der Thusen, J. H. & van Paassen, J. A systematic review of pathological findings in COVID-19: a pathophysiological timeline and possible mechanisms of disease progression. Mod. Pathol. 33, 2128–2138, doi:10.1038/s41379-020-0603-3 (2020).

13 Liu, Q. et al. Lung regeneration by multipotent stem cells residing at the bronchioalveolar-duct junction. Nat Genet 51, 728–738, doi:10.1038/s41588-019-0346-6 (2019).

14 Liu, K. et al. Bi-directional differentiation of single bronchioalveolar stem cells during lung repair. Cell Discov 6, 1, doi:10.1038/s41421-019-0132-8 (2020).

15 Kathiriya, J. J., Brumwell, A. N., Jackson, J. R., Tang, X. & Chapman, H. A. Distinct Airway Epithelial Stem Cells Hide among Club Cells but Mobilize to Promote Alveolar Regeneration. Cell Stem Cell 26, 346–358 e344, doi:10.1016/j.stem.2019.12.014 (2020).

16 Barkauskas, C. E. A Specialized Few Among Many: Identification of a Novel Lung Epithelial Stem Cell Population. Cell Stem Cell 26, 295–296, doi:10.1016/j.stem.2020.02.010 (2020).

17 Vaughan, A. E. et al. Lineage-negative progenitors mobilize to regenerate lung epithelium after major injury. Nature 517, 621–625, doi:10.1038/nature14112 (2015).

18 Jiang, P. et al. Ineffectual Type 2-to-Type 1 Alveolar Epithelial Cell Differentiation in Idiopathic Pulmonary Fibrosis: Persistence of the KRT8(hi) Transitional State. Am J Respir Crit Care Med 201, 1443–1447, doi:10.1164/rccm.201909-1726LE (2020).

19 Strunz, M. et al. Alveolar regeneration through a Krt8+ transitional stem cell state that persists in human lung fibrosis. Nat Commun 11, 3559, doi:10.1038/s41467-020-17358-3 (2020).

20 Riemondy, K. A. et al. Single cell RNA sequencing identifies TGFβ as a key regenerative cue following LPS-induced lung injury. JCI Insight 5, doi:10.1172/jci.insight.123637 (2019).

21 Kobayashi, Y. et al. Persistence of a regeneration-associated, transitional alveolar epithelial cell state in pulmonary fibrosis. Nat. Cell Biol. 22, 934–946, doi:10.1038/s41556-020-0542-8 (2020).

22 Jansing, N. L. et al. Unbiased Quantitation of Alveolar Type II to Alveolar Type I Cell Transdifferentiation during Repair after Lung Injury in Mice. Am J Respir Cell Mol Biol 57, 519–526, doi:10.1165/rcmb.2017-0037MA (2017).

23 Wu, H. et al. Progressive Pulmonary Fibrosis Is Caused by Elevated Mechanical Tension on Alveolar Stem Cells. Cell 180, 107-121.e117, doi:10.1016/j.cell.2019.11.027 (2020).

24 Delorey, T. M. et al. COVID-19 tissue atlases reveal SARS-CoV-2 pathology and cellular targets. Nature, doi:10.1038/s41586-021-03570-8 (2021).

25 Bharat, A. et al. Lung transplantation for patients with severe COVID-19. Sci Transl Med 12, doi:10.1126/scitranslmed.abe4282 (2020).

26 Chan, J. F. et al. Simulation of the Clinical and Pathological Manifestations of Coronavirus Disease 2019 (COVID-19) in a Golden Syrian Hamster Model: Implications for Disease Pathogenesis and Transmissibility. Clin. Infect. Dis. 71, 2428–2446, doi:10.1093/cid/ciaa325 (2020).

27 Sia, S. F. et al. Pathogenesis and transmission of SARS-CoV-2 in golden hamsters. Nature 583, 834–838, doi:10.1038/s41586-020-2342-5 (2020).

28 Muñoz-Fontela, C. et al. Animal models for COVID-19. Nature 586, 509–515, doi:10.1038/s41586-020-2787-6 (2020).

29 Ciurkiewicz, M. et al. Ferrets are valuable models for SARS-CoV-2 research. Vet Pathol 59, 661–672, doi:10.1177/03009858211071012 (2022).

30 Winkler, E. S. et al. SARS-CoV-2 infection of human ACE2-transgenic mice causes severe lung inflammation and impaired function. Nat Immunol 21, 1327–1335, doi:10.1038/s41590-020-0778-2 (2020).

31 Becker, K. et al. Vasculitis and Neutrophil Extracellular Traps in Lungs of Golden Syrian Hamsters With SARS-CoV-2. Front. Immunol. 12, 640842, doi:10.3389/fimmu.2021.640842 (2021).

32 Allnoch, L. et al. Vascular Inflammation Is Associated with Loss of Aquaporin 1 Expression on Endothelial Cells and Increased Fluid Leakage in SARS-CoV-2 Infected Golden Syrian Hamsters. Viruses 13, doi:10.3390/v13040639 (2021).

33 Mulka, K. R. et al. Progression and Resolution of Severe Acute Respiratory Syndrome Coronavirus 2 (SARS-CoV-2) Infection in Golden Syrian Hamsters. Am J Pathol 192, 195–207, doi:10.1016/j.ajpath.2021.10.009 (2022).

34 Armando, F. et al. SARS-CoV-2 Omicron variant causes mild pathology in the upper and lower respiratory tract of hamsters. Nat Commun 13, 3519, doi:10.1038/s41467-022-31200-y (2022).

35 Lantz, R. C., Birch, K., Hinton, D. E. & Burrell, R. Morphometric changes of the lung induced by inhaled bacterial endotoxin. Exp Mol Pathol 43, 305–320, doi:10.1016/0014-4800(85)90068-1 (1985).

36 Ochs, M. et al. Using electron microscopes to look into the lung. Histochem Cell Biol 146, 695–707, doi:10.1007/s00418-016-1502-z (2016).

37 Chen, J., Wu, H., Yu, Y. & Tang, N. Pulmonary alveolar regeneration in adult COVID-19 patients. Cell Res. 30, 708–710, doi:10.1038/s41422-020-0369-7 (2020).

38 Zuo, W. et al. p63(+)Krt5(+) distal airway stem cells are essential for lung regeneration. Nature 517, 616–620, doi:10.1038/nature13903 (2015).

39 Zacharias, W. J. et al. Regeneration of the lung alveolus by an evolutionarily conserved epithelial progenitor. Nature 555, 251–255, doi:10.1038/nature25786 (2018).

40 Parekh, K. R. et al. Stem cells and lung regeneration. Am. J. Physiol. Cell Physiol. 319, C675–C693, doi:10.1152/ajpcell.00036.2020 (2020).

41 Smirnova, N. F. et al. Detection and quantification of epithelial progenitor cell populations in human healthy and IPF lungs. Respir. Res. 17, 83, doi:10.1186/s12931-016-0404-x (2016).

42 Musah, S., Chen, J. & Hoyle, G. W. Repair of tracheal epithelium by basal cells after chlorine-induced injury. Respir. Res. 13, 107, doi:10.1186/1465-9921-13-107 (2012).

43 Nouailles, G. et al. Temporal omics analysis in Syrian hamsters unravel cellular effector responses to moderate COVID-19. Nat Commun 12, 4869, doi:10.1038/s41467-021-25030-7 (2021).

44 Xie, T. et al. Abnormal respiratory progenitors in fibrotic lung injury. Stem Cell. Res. Ther. 13, 64, doi:10.1186/s13287-022-02737-y (2022).

45 Carvallo, F. R. & Stevenson, V. B. Interstitial pneumonia and diffuse alveolar damage in domestic animals. Vet. Pathol. 59, 586–601, doi:10.1177/03009858221082228 (2022).

46 Guler, S. A. et al. Pulmonary function and radiological features 4 months after COVID-19: first results from the national prospective observational Swiss COVID-19 lung study. Eur. Respir. J. 57, doi:10.1183/13993003.03690-2020 (2021).

47 Desai, A. D., Lavelle, M., Boursiquot, B. C. & Wan, E. Y. Long-term complications of COVID-19. Am. J. Physiol. Cell Physiol. 322, C1–c11, doi:10.1152/ajpcell.00375.2021 (2022).

48 Singh, I. & Joseph, P. Short and Long Term Non-Invasive Cardiopulmonary Exercise Assessment in previously Hospitalized COVID-19 Patients. Eur. Respir. J., doi:10.1183/13993003.01739-2022 (2022).

49 Frere, J. J. et al. SARS-CoV-2 infection in hamsters and humans results in lasting and unique systemic perturbations post recovery. Sci Transl Med, eabq3059, doi:10.1126/scitranslmed.abq3059 (2022).

50 Baptiste, N. & Prives, C. p53 in the cytoplasm: a question of overkill? Cell 116, 487–489, doi:10.1016/s0092-8674(04)00164-3 (2004).

51 McGregor, A. L., Hsia, C. R. & Lammerding, J. Squish and squeeze-the nucleus as a physical barrier during migration in confined environments. Curr. Opin. Cell Biol. 40, 32–40, doi:10.1016/j.ceb.2016.01.011 (2016).

52 Dinnon, K. H., 3rd et al. SARS-CoV-2 infection produces chronic pulmonary epithelial and immune cell dysfunction with fibrosis in mice. Sci. Transl. Med. 14, eabo5070, doi:10.1126/scitranslmed.abo5070 (2022).

53 Yao, C. et al. Senescence of Alveolar Type 2 Cells Drives Progressive Pulmonary Fibrosis. Am. J. Respir. Crit. Care Med. 203, 707–717, doi:10.1164/rccm.202004-1274OC (2021).

54 Lehmann, M. et al. Chronic WNT/β-catenin signaling induces cellular senescence in lung epithelial cells. Cell. Signal. 70, 109588, doi:10.1016/j.cellsig.2020.109588 (2020).

55 Finn, J. et al. Dlk1-Mediated Temporal Regulation of Notch Signaling Is Required for Differentiation of Alveolar Type II to Type I Cells during Repair. Cell Rep. 26, 2942-2954.e2945, doi:10.1016/j.celrep.2019.02.046 (2019).

56 Lee, S. et al. Virus-induced senescence is a driver and therapeutic target in COVID-19. Nature 599, 283–289, doi:10.1038/s41586-021-03995-1 (2021).

57 Rosenke, K. et al. Defining the Syrian hamster as a highly susceptible preclinical model for SARS-CoV-2 infection. Emerg Microbes Infect 9, 2673–2684, doi:10.1080/22221751.2020.1858177 (2020).

58 Song, Z. et al. SARS-CoV-2 Causes a Systemically Multiple Organs Damages and Dissemination in Hamsters. Front. Microbiol. 11, 618891, doi:10.3389/fmicb.2020.618891 (2020).

59 Hinz, B. Mechanical aspects of lung fibrosis: a spotlight on the myofibroblast. Proc. Am. Thorac. Soc. 9, 137–147, doi:10.1513/pats.201202-017AW (2012).

60 Braga, T. T., Agudelo, J. S. & Camara, N. O. Macrophages During the Fibrotic Process: M2 as Friend and Foe. Front. Immunol. 6, 602, doi:10.3389/fimmu.2015.00602 (2015).

61 Hill, C., Jones, M. G., Davies, D. E. & Wang, Y. Epithelial-mesenchymal transition contributes to pulmonary fibrosis via aberrant epithelial/fibroblastic cross-talk. J Lung Health Dis 3, 31–35 (2019).

62 Yao, L. et al. Paracrine signalling during ZEB1-mediated epithelial-mesenchymal transition augments local myofibroblast differentiation in lung fibrosis. Cell Death Differ. 26, 943–957, doi:10.1038/s41418-018-0175-7 (2019).

63 Ackermann, M. et al. The fatal trajectory of pulmonary COVID-19 is driven by lobular ischemia and fibrotic remodelling. EBioMedicine 85, 104296, doi:10.1016/j.ebiom.2022.104296 (2022).

64 Zhao, Z. et al. Single-cell analysis identified lung progenitor cells in COVID-19 patients. Cell Prolif 53, e12931, doi:10.1111/cpr.12931 (2020).

65 Schreiner, T. et al. SARS-CoV-2 Infection Dysregulates Cilia and Basal Cell Homeostasis in the Respiratory Epithelium of Hamsters. Int. J. Mol. Sci. 23, doi:10.3390/ijms23095124 (2022).

66 Armando, F. et al. Intratumoral Canine Distemper Virus Infection Inhibits Tumor Growth by Modulation of the Tumor Microenvironment in a Murine Xenograft Model of Canine Histiocytic Sarcoma. Int. J. Mol. Sci. 22, doi:10.3390/ijms22073578 (2021).

67 Armando, F. et al. Mesenchymal to epithelial transition driven by canine distemper virus infection of canine histiocytic sarcoma cells contributes to a reduced cell motility in vitro. J. Cell. Mol. Med. 24, 9332–9348, doi:10.1111/jcmm.15585 (2020).

68 Armando, F. et al. Endocanalicular transendothelial crossing (ETC): A novel intravasation mode used by HEK-EBNA293-VEGF-D cells during the metastatic process in a xenograft model. PLoS One 15, e0239932, doi:10.1371/journal.pone.0239932 (2020).

69 Bankhead, P. et al. QuPath: Open source software for digital pathology image analysis. Sci. Rep. 7, 16878, doi:10.1038/s41598-017-17204-5 (2017).

70 R_Core_Team. R: A language and environment for statistical computing. R Foundation for Statistical Computing, Vienna, Austria URL http://www.R-project.org/ (2014).

71 Satija, R., Farrell, J. A., Gennert, D., Schier, A. F. & Regev, A. Spatial reconstruction of single-cell gene expression data. Nat. Biotechnol. 33, 495–502, doi:10.1038/nbt.3192 (2015).

72 Butler, A., Hoffman, P., Smibert, P., Papalexi, E. & Satija, R. Integrating single-cell transcriptomic data across different conditions, technologies, and species. Nat. Biotechnol. 36, 411–420, doi:10.1038/nbt.4096 (2018).

73 Stuart, T. et al. Comprehensive Integration of Single-Cell Data. Cell 177, 1888-1902.e1821, doi:10.1016/j.cell.2019.05.031 (2019).

74 Hao, Y. et al. Integrated analysis of multimodal single-cell data. Cell 184, 3573-3587.e3529, doi:10.1016/j.cell.2021.04.048 (2021).

75 Liberzon, A. et al. The Molecular Signatures Database (MSigDB) hallmark gene set collection. Cell Syst 1, 417–425, doi:10.1016/j.cels.2015.12.004 (2015).

